# Mechanical confinement and DDR1 signalling synergise to regulate collagen-induced apoptosis in rhabdomyosarcoma cells

**DOI:** 10.1101/2022.04.26.489526

**Authors:** Jordi Gonzalez-Molina, Katharina Miria Kirchhof, Bhavik Rathod, Lidia Moyano-Galceran, Maria Calvo-Noriega, Georgia Kokaraki, Astrid Bjørkøy, Monika Ehnman, Joseph W. Carlson, Kaisa Lehti

**Affiliations:** Department of Microbiology, Tumor and Cell Biology, Karolinska Institutet, Stockholm, Sweden; Department of Oncology-Pathology, Karolinska Institutet, Stockholm, Sweden; Department of Laboratory Medicine, Division of Pathology, Karolinska Institutet, Stockholm, Sweden; Keck School of Medicine, University of Southern California, Los Angeles, US; Department of Physics, Norwegian University of Science and Technology, Trondheim, Norway; Department of Biomedical Laboratory Science, Norwegian University of Science and Technology, Trondheim, Norway

## Abstract

Fibrillar collagens promote cell proliferation, migration, and survival in various epithelial cancers and are generally associated with tumour aggressiveness. However, the impact of fibrillar collagens on soft tissue sarcoma behaviour remains poorly understood.

Unexpectedly, we find here that fibrillar collagen-related gene expression is associated with favourable patient prognosis in rhabdomyosarcoma. By developing and using collagen matrices with distinct stiffness and *in vivo*-like microarchitectures, we uncover that the activation of DDR1 has pro-apoptotic and integrin β1 pro-survival function, specifically in 3D rhabdomyosarcoma cell cultures. We demonstrate that rhabdomyosarcoma cell-intrinsic or extrinsic matrix remodelling promotes cell survival. Mechanistically, we find that the 3D-specific collagen-induced apoptosis results from a dual DDR1-independent and a synergistic DDR1-dependent TRPV4-mediated response to mechanical confinement. Altogether, our results indicate that dense microfibrillar collagen-rich microenvironments are detrimental to rhabdomyosarcoma cells through an apoptotic response orchestrated by the induction of DDR1 signalling and mechanical confinement. This mechanism helps to explain the preference of rhabdomyosarcoma cells to grow in and metastasise to low fibrillar collagen microenvironments such as the lung.

## INTRODUCTION

Sarcomas constitute approximately 7% of all paediatric cancers, with rhabdomyosarcoma (RMS) being the most common type (*1*). Despite their rareness, sarcomas are estimated to cause 13% of cancer-related deaths in children (*2*). Current multimodal treatments have been successful to improve the 5-year event-free survival rate of patients with localised disease to at least 70%. Yet, the survival of patients with metastatic or relapse disease drops below 30% (*2*). Understanding the distinct factors regulating RMS tumourigenesis and disease progression will be fundamental to avoid disease recurrence and improve metastasis treatment.

Both cell-intrinsic characteristics and a favourable tumour microenvironment (TME) are necessary for cancer cell survival, proliferation, and the formation of primary solid tumours and metastases (*3*). A central component of the TME-cell communication is the extracellular matrix (ECM), a 3D network of diverse macromolecules that provides cells with a physical support and biochemical cues. Cells adhere to the ECM molecules through transmembrane proteins such as integrins, discoidin domain receptors (DDRs), syndecans, and CD44, which trigger the activation of downstream signalling pathways upon ligand binding. These pathways control fundamental processes in cancer development and progression such as cell proliferation, survival, and migration depending on both the expressed receptor repertoire and the ECM composition (*4, 5*). Moreover, both the composition and mechanical properties of the ECM, highly dependent especially on the expression and crosslinking of fibrillar collagens, are known to affect disease aggressiveness, drug delivery, and induce therapy resistance in various carcinomas (*6*). Instead, on soft tissue sarcomas, our current knowledge of the impact of fibrillar collagens remains limited. Considering the biological diversity, with more than 80 soft tissue sarcoma types reported, the effects of collagenous ECM in these neoplasms may be heterogeneous (*7*).

Dense and stiff ECMs impose physical constraints that limit tumour growth and cell invasion, but are often associated with tumour aggressiveness (*8*). Enzymatic ECM degradation, mainly through metalloproteinases such as matrix metalloproteinases (MMPs), and cell-induced tensile forces allow cells to overcome these physical limitations and form tumours or colonise other tissues. Thus, ECM remodelling activity is commonly increased in cancer cells (*9*). Mesenchymal cells typically present high ECM remodelling capabilities. However, how these characteristics are not necessarily conserved or are altered in different types of sarcoma cells, which are of mesenchymal origin, is unclear. For instance, RMS tumours generally express low MMP14, the membrane-type MMP with central functions in collagen degradation (*10*). The cell responses to the inability to remodel or degrade the surrounding dense ECM are also elusive, but have been linked to a reversible quiescent state known as cell dormancy, which is associated with therapy resistance and disease relapse (*11*). Hence, understanding the relationship between tumour cell-intrinsic and ECM characteristics in these processes is fundamental to predict cell behaviour *in vivo* and to improve therapy response, which is of critical importance for aggressive soft tissue sarcomas.

Here, we show how expression of fibrillar collagen genes and collagen biosynthesis genes are associated with favourable prognosis in RMS. Using *in vitro* 2D and 3D collagen I-based models with varying fibre microarchitecture and stiffness, we show that soft 3D microfibrillar collagen induces an apoptotic response in RMS cells. We determine that 3D microfibrillar collagen and collagen-mediated mechanical confinement regulate DDR1 signalling to induce apoptosis. In contrast, we find that, in specific cell lines, stiffer bundled collagen promotes cell survival through the activation of integrin β1. Moreover, we show that the 3D-specific apoptotic response is caused by DDR1-dependent and -independent mechanisms of mechanical confinement, which can be overcome by ECM remodelling. Altogether, these results show that microenvironmental factors imposed by collagen coupled with cell-intrinsic characteristics such as the expression of DDR1 and the ability to remodel fibrillar collagen matrices determine RMS cell survival and proliferation.

## RESULTS

### Expression of fibrillar collagen-related genes is associated with favourable prognosis in rhabdomyosarcoma

To explore the impact of fibrillar collagen on sarcoma aggressiveness, we evaluated the association between gene expression of fibrillar collagen or collagen biosynthesis/crosslinking-associated genes and sarcoma patient prognosis in three distinct sarcoma cohorts (*12*). In The Cancer Genome Atlas Sarcoma cohort (TCGA-SARC), which includes 262 cases of diverse types of adult soft tissue sarcomas, high expression of 4 out of 11 fibrillar collagen (FibrillarCOL) genes and 5/14 collagen biosynthesis (COLbio) genes was associated with unfavourable prognosis, whilst only one COLbio gene, FMOD, was associated with better patient survival (Fig. 1A and Supplementary Table 1). In the collagen-rich osteosarcomas (GSE21257 cohort, n = 53), high expression of a FibrillarCOL (1/11) and a COLbio (1/14) genes, was likewise associated with poor prognosis. In contrast, in rhabdomyosarcoma (RMS) tumours (ITCC cohort, n=101), the expression of FibrillarCOL (2/11) and COLbio (4/14) genes was associated specifically with favourable prognosis.

**Figure 1.**
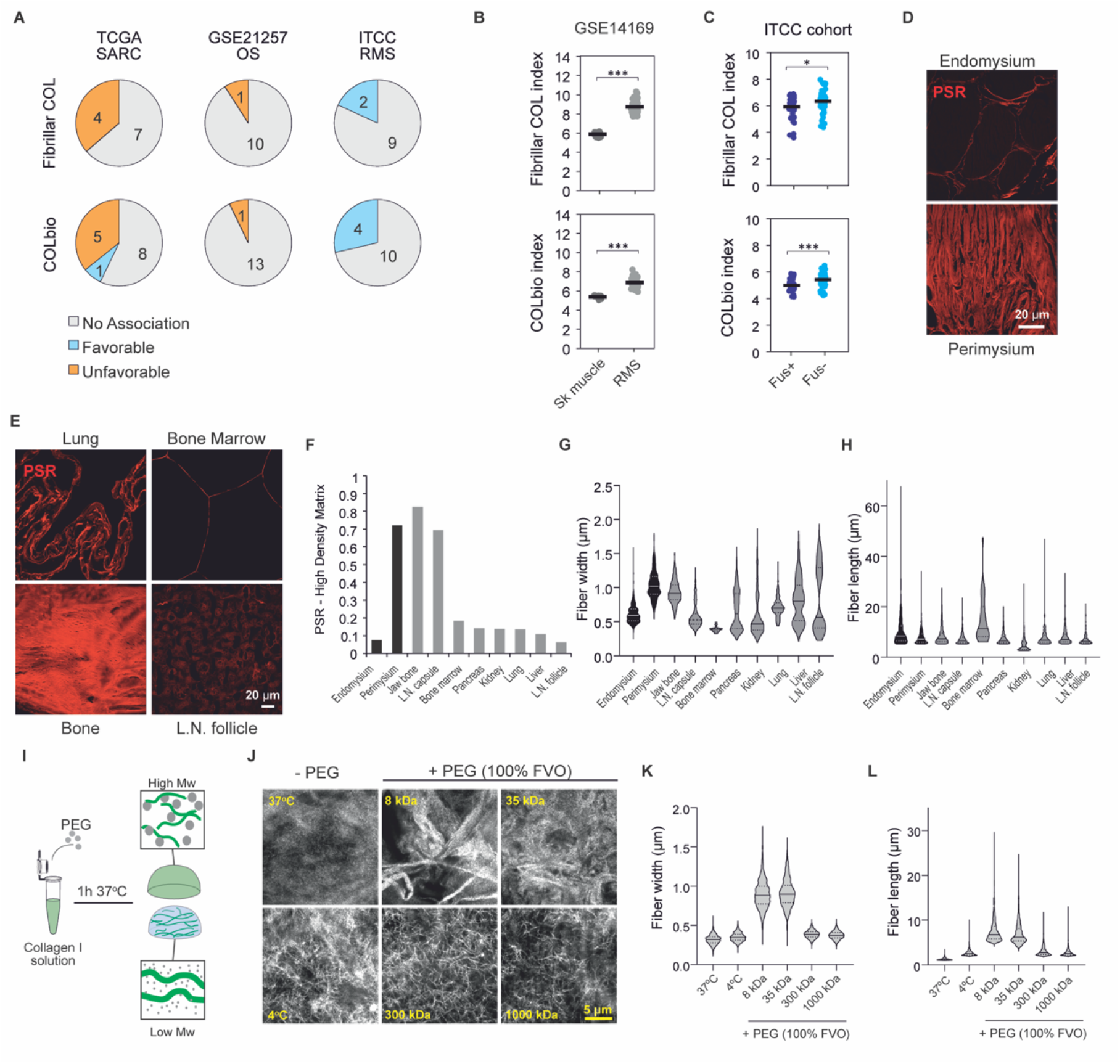
Tissue fibrillar collagen analysis in rhabdomyosarcoma and common metastatic sites and tissue collagen-mimicking matrices for *in vitro* studies. **A**, Association between fibrillar collagen and collagen biosynthesis/crosslinking gene expression and patient survival from the TCGA-SARC (n=262), GSE21257 (n=53), and ITCC (n=101) cohorts. **B, C**, Average gene expression of all fibrillar collagens (top) and collagen biosynthesis and crosslinking genes (bottom) shows increased expression in rhabdomyosarcoma tissues (RMS) compared to skeletal muscle tissue (sk muscle) (**B**) and higher expression in fusion-negative RMS (Fus-) than in fusion-positive RMS (Fus+) (**C**). Bars indicate the average expression. **D**, Picrosirius red (PSR) staining shows fibrillar collagen content and structure in the connective tissues of human skeletal muscle. Scale bar indicates 20 μm. **E**, PSR staining shows diverse structure and content of fibrillar collagens in common tissue sites of RMS metastasis. Scale bar indicates 20 μm. **F**, Quantification of PSR-high density area shows diverse density in skeletal muscle tissue and common sites of RMS metastasis (n ≥ 3 10x magnification fields per tissue). **G**, Quantification of fibre width based on confocal images of PSR-stained tissues show heterogeneity between RMS-relevant tissues (n ≥ 33 fibres per tissue). **H**, Quantification of fibre length (within the limits of the image) based on confocal images of PSR-stained tissues shows heterogeneity between RMS-relevant tissues (n ≥ 33 fibres per tissue). **I**, Schematic illustration of the method for modifying collagen microarchitecture with PEG-induce macromolecular crowding. **J**, Confocal reflectance microscopy images of collagen hydrogels where fibrillogenesis was performed without PEG or with PEG at distinct molecular weights at a 100% fraction volume of occupancy (FVO). Scale bar indicates 5 μm. **K**,**L**, Fibre width (**K**) and length (**L**) quantification of the collagen hydrogels in (**J**). Violin plots represent the median (solid line) and the interquartile range (dotted lines). *** p < 0.001, * p < 0.05.

To explore further the dysregulation of fibrillar collagen gene expression in RMS, we compared the average expression of FibrillarCOL and COLbio genes between RMS tumours and normal skeletal muscle tissue (GSE14169). As seen in various other tumour types (*13*), RMS tumours show enhanced expression of both FibrillarCOL and COLbio genes compared to the normal tissue (Fig. 1B). Finally, to investigate the differences between RMS molecular subtypes, which have distinct clinical outcomes (*14, 15*), we compared the expression of fibrillar collagen-related genes between fusion-positive and fusion-negative tumours. Of note, fusion-positive tumours, typically associated with worse prognosis, showed significantly lower average FibrillarCOL and COLbio gene expression than fusion-negative tumours (Fig. 1C). Altogether, these data indicate that, opposite to various previously reported carcinoma types as well as the other sarcomas analysed here, fibrillar collagen-related gene expression is unexpectedly linked to less aggressive RMS tumours.

### Tissue-like collagen structures can be modelled via macromolecular crowding in 3D collagen

Recreating the microarchitecture and mechanics of native fibrillar collagens *in vitro* is fundamental to understand their impact on cell behaviour (*16, 17*). Considering that the structural collagen properties in tissues prior to malignant tissue remodelling can determine the ability of cancer cells to form tumours, we stained tissue collagen with picrosirius red in skeletal muscle, as well as lung, lymph node, bone marrow, bone, pancreas, kidney, and liver tissues representing the reported RMS metastatic sites (*18*). Both skeletal muscle and the metastasis-permissive tissues generally showed an overall low density of fibrillar collagen, with the exception of the specialized connective tissues in skeletal muscle (perimysium) and lymph node (capsule), as well as bone (Fig. 1D-F). The collagen fibre width and length showed high inter-tissue variability, with tissues such as the perimysium showing thick (>1 μm) and long (up to >60 μm; likely an underestimation, as longer fibre measurements were limited by the image field) fibres, and bone marrow only presenting thin (<0.5 μm) and short (up to 20 μm) microfibres (Fig. 1G,H).

Given this observed heterogeneity in collagen characteristics, reliable models used to study the response of RMS cells to collagen should typify these structural differences. We hypothesised that induction of macromolecular crowding, known to alter collagen microarchitecture (*19*), during the collagen fibrillogenesis process would lead to the formation of the variable tissue-like collagen fibres. We used polyethylene glycol (PEG) as the macromolecular crowder because it is inert and available at defined molecular weights, allowing the control of the viscosity as well as the volume occupied by the PEG molecules in the solution. We also considered that, at high fraction volume of occupancy (FVO), the length and width of the formed collagen type I fibres increases. Enhancing the viscosity of the solution, by varying the PEG molecular weight, would instead decrease the collagen fibre length and thickness (Fig. 1I). Indeed, at 100% FVO, the less viscous 8 kDa PEG solution induced the formation of thick and long collagen I fibres, or bundles, of comparable width and length to those observed in perimysium, bone, or lung tissues, whereas 1,000 kDa PEG supported the formation of shorter fibres, comparable to the microfibres in kidney tissue (Fig. 1J-L). The middle molecular weight PEGs (35-300 kDa) in turn, produced intermediate sized fibres. In the absence of macromolecular crowding, microfibres were even thinner and shorter than with 1,000 kDa PEG, comparable to bone marrow. Fibrillogenesis at 4ºC, also reported to increase fibre width and length (*17*), did not achieve similar collagen structures as those induced by the 8 kDa PEG. These results indicate that macromolecular crowding is a useful method to form relevant tissue-like collagen models to study RMS, including the large collagen bundles and microfibrillar collagen structures mimicking lung/bone/perimysium and bone marrow, respectively.

### Collagen microarchitecture heterogeneously affects rhabdomyosarcoma cell proliferation, adhesion, and mechanotransduction

To investigate the response of RMS cells to distinct tissue-like collagen structures, we seeded six different RMS cell lines (including fusion+ and fusion-; Supplementary Table 2) on top of microfibrillar collagen (MF-collagen; formed at low macromolecular crowding) or bundled collagen (B-collagen; induced by 8 kDa PEG) matrices. For comparison, we used 4 osteosarcoma (OS) cell lines, derived from and producing collagen-rich microenvironments (Supplementary Fig. 1A,B). Proliferation rates assessed by nuclear Ki67 positivity showed that, unlike in OS cells, the response of RMS cells to collagen was dependent on collagen microarchitecture, with the cell lines RUCH2 and RD showing a trend to higher proliferation and the cell line RMS a lower proliferation on B-collagen (Fig. 2A-C). Whilst integrin β1 activation and YAP/TAZ nuclear localisation, both linked to cell proliferation (*20*), increased in B-collagen compared to MF-collagen, the levels of activation were generally lower in RMS than in OS (Fig. 2D-I). As an exception amongst the RMS cell lines, the RUCH2 cell line presented the highest total expression of fully glycosylated, mature integrin β1 (Supplementary Fig. 1C) (*21*), in conjunction with relatively high integrin β1 activation and YAP/TAZ nuclear localisation in B-collagen at comparable levels to the OS cell lines. Integrin β1 activation and YAP/TAZ nuclear localisation are generally promoted in stiff matrices (*20*). Indeed, B-collagen matrices were significantly stiffer than MF-collagen matrices (Young ‘s modulus of 1.62 kPa for B-collagen vs 0.32 kPa for MF-collagen; p=0.00011), as characterised by atomic force microscopy (Supplementary Fig. 1D). Similarly, adhesion to supraphysiologically stiff collagen-coated glass did not have a major impact on proliferation (Fig. 2C), but induced relatively high integrin β1 activation and YAP/TAZ nuclear:cytoplasmic ratio in RMS cells (Fig. 2F,I).These results indicate that RMS cell proliferation, adhesion, and mechanotransduction are sensitive to collagen microarchitecture, tightly linked to matrix stiffness. However, integrin β1 activation and YAP/TAZ nuclear localisation alone do not explain the observed variability between cell lines, suggesting that other factors influence RMS cell proliferation on collagen matrices.

**Figure 2.**
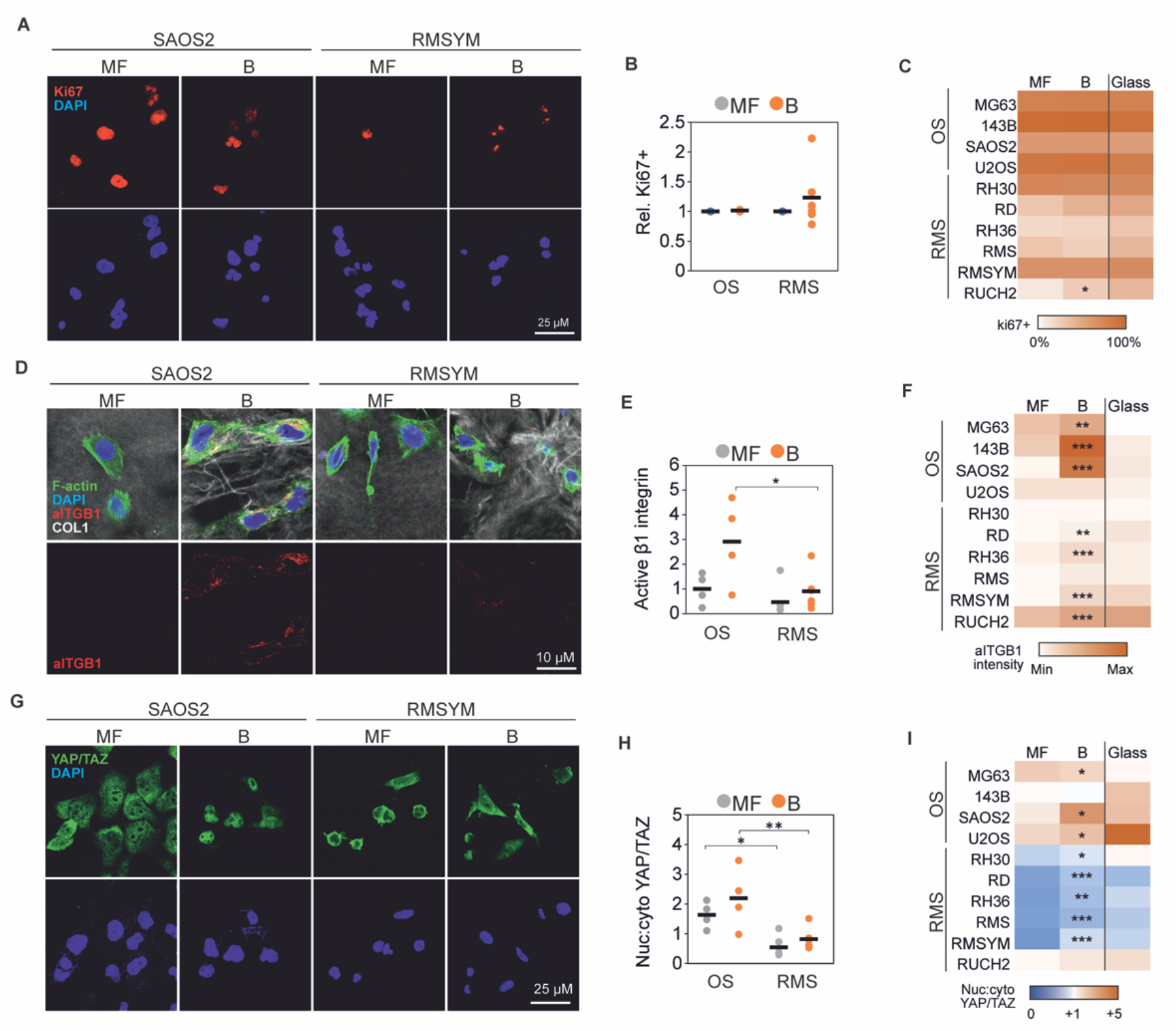
Collagen microarchitecture regulates rhabdomyosarcoma cell proliferation. **A**, Representative examples of Ki67 staining of an osteosarcoma (SAOS2) and a rhabdomyosarcoma (RMSYM) cell line seeded on MF-or B-collagen. Scale bar indicates 25 μm. **B**, Quantification of the percentage of Ki67+ cells relative to the percentage of Ki67+ cells on MF-collagen showing heterogeneous response in rhabdomyosarcoma cell lines. **C**, Percentage of Ki67+ cells in each osteosarcoma (OS) and rhabdomyosarcoma (RMS) cell line on MF-collagen, B-collagen, or on collagen-coated glass. **D**, Representative examples of SAOS2 and RMSYM cells showing increased active integrin β1 (aITGB1) staining in B-collagen conditions. Scale bar represents 10 μm. **E**, Quantification of active integrin β1 per cell normalised to the average of all OS cell lines on MF-collagen. **F**, Quantification of active integrin β1 per cell line on MF-collagen, B-collagen, or collagen-coated glass. **G**, Representative examples of SAOS2 and RMSYM cells showing distinct subcellular localization of YAP/TAZ. Scale bar represents 25 μm. **H, I**, Quantification of nuclear:cytoplasmic ratio of YAP/TAZ in OS and RMS cell lines showing generally lower nuclear localization in RMS cell lines. Each data point represents the average of 3 independent experiments of one cell line. Bars indicate the average of all cell lines. *** p < 0.001, ** p < 0.01, * p < 0.05 (indicating comparison between MF and B in (**C, F, I**).

### Collagen induces an apoptotic response on RMS cells dependent on dimensionality and microarchitecuture

Next, to simulate the 3D confining effect of the *in vivo* TME, we embedded the distinct RMS and OS cells within 3D MF- and B-collagen matrices. Similarly to the 2D cultures on top of matrices, the RMS cells showed a cell line-dependent proliferative response to the alteration of collagen microarchitecture after 24 h, whereas the proliferation of OS cells remained unaffected also in 3D (Fig. 3A). After 4-days in culture, the cell lines RH36, RMS, and RMSYM were still barely detectable in MF-collagen (Fig. 3B). In B-collagen, RMSYM cells instead formed cell spheroids/clusters, indicative of cell proliferation (Fig. 3B). This growth suppressive effect of MF-collagen was not seen in RD, RH30, or RUCH2 or in any of the tested OS cell lines (Fig. 3B and Supplementary Fig. 2A). To investigate whether the proliferation trends observed in 2D and 3D conditions were similar, we compared the ratio of Ki67+ cells between 2D and 3D in MF- and B-collagen. The ratio of Ki67+ OS cells was constant, indicating no variations in proliferation in any of the conditions (Fig. 3C). However, RMS cell lines showed heterogeneous responses to both dimensionality and collagen microarchitecture and presented generally lower proliferation in 3D B-collagen (Fig. 3C). This lower proliferation in 3D B-collagen was observed in RH30, RD, RH36, RMS, and RMSYM (Fig. 3D). Instead, RUCH2 showed higher proliferation in 3D compared to 2D, further indicating that this cell line presents characteristics that are not shared with the other cell lines (Fig. 3D). To investigate whether the low proliferation and low cell detection after 4 days in culture was the result of an apoptotic response triggered by collagen, we stained cleaved-caspase 3 (cl-caspase 3) in cells after 24 h in culture (Fig. 3E). In RH36, RMS, and RMSYM, apoptosis was considerably enhanced (11-60% cl-caspase 3+) in 3D collagen, and in RD at lower levels (Fig. 3F). Of note, the increase in cl-caspase 3+ RMSYM cells was only observed in 3D MF-collagen. However, these cell lines were able to proliferate in non-collagen 3D matrices such as 0.2% agarose and fibrin (Supplementary Fig. 2B). Although the effect of distinct microarchitecture and mechanics in these alternative matrices likely also affect the cells, these results altogether suggest that MF- and B-collagens can induce differential growth-suppressive and apoptotic responses particularly in RMS cells.

**Figure 3.**
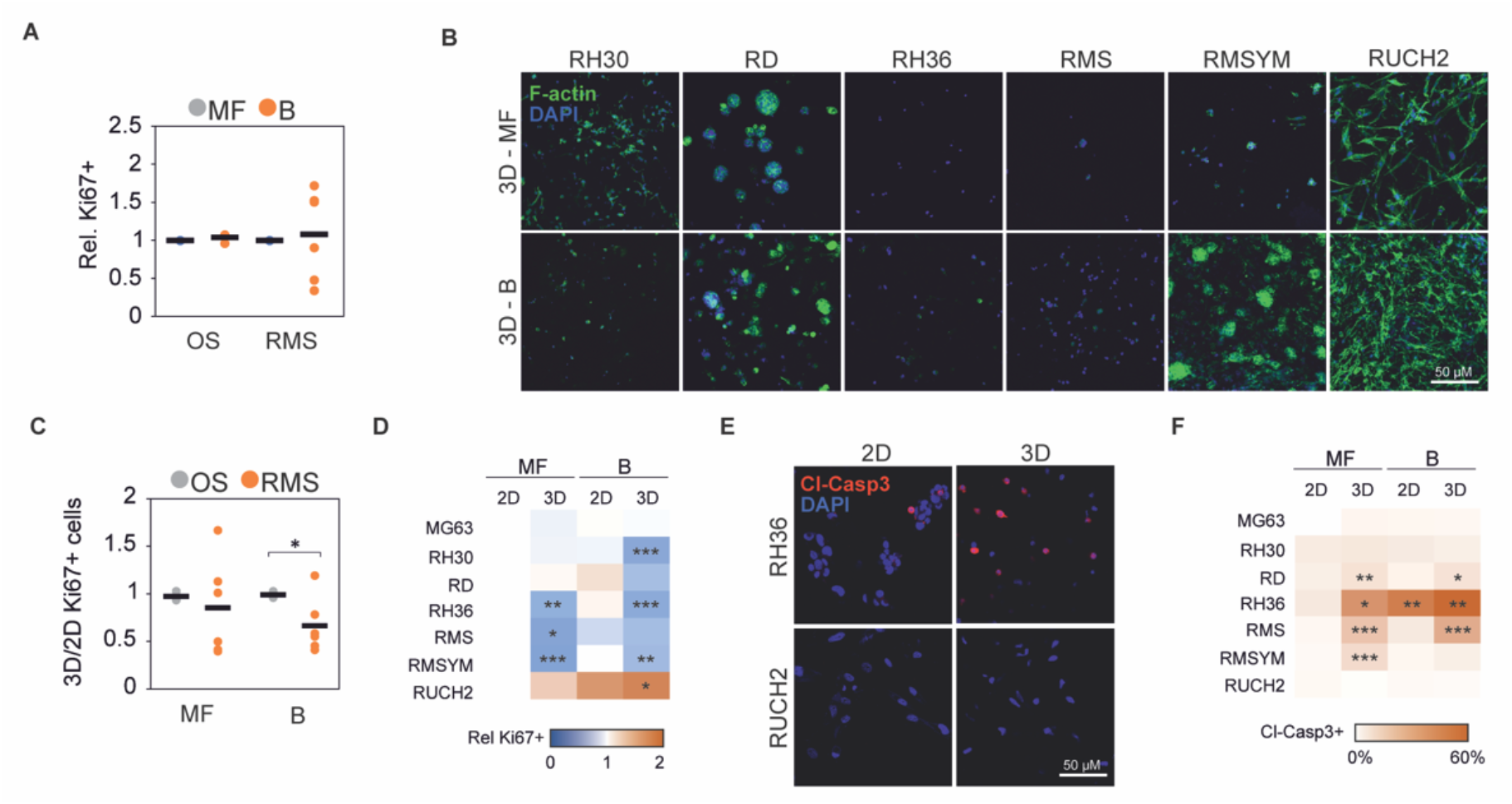
3D collagen culture induces a microarchitecture-dependent apoptotic response on rhabdomyosarcoma cells. **A**, Quantification of the percentage of Ki67+ cells relative to the percentage of Ki67+ cells in 3D MF-collagen 24 h after embedding within 3D collagen matrices. **B**, Representative examples of phalloidin-stained rhabdomyosarcoma cell lines grown in 3D MF- or B-collagen for 4 days. Scale bar indicates 50 μm. **C**, Ratio of percentage of Ki67+ cells between 24 h 3D and 2D cultures in B- or MF-collagen showing that RMS cell lines response to collagen microarchitecture and dimensionality is cell line-dependent. **D**, Heatmap representing the relative number of Ki67+ cells at distinct dimensionality and collagen microarchitecture in each RMS cells line and in the OS cell line MG63 used for comparison. **E**, Representative images of cleaved-caspase 3 (Cl-Casp3)-stained RH36 and RUCH2 cells cultured for 24 h in 2D or 3D MF- collagen. **F**, Percentage of Cl-Caps3+ cells per cell line cultured for 24 h in the indicated conditions. Each data point represents the average of 3 independent experiments of one cell line. Bars indicate the average of all cell lines. *** p < 0.001, ** p < 0.01, * p < 0.05.

To investigate whether such apoptotic responses were cell cycle- and YAP/TAZ-dependent, we used L-mimosine or S-Trityl-L-Cysteine (STLC) to block cell cycle progression or verteporfin to inhibit YAP/TAZ. Blocking cell cycle in apoptotic RH36, RMS, and RMSYM cell lines or inhibiting YAP/TAZ in the distinct OS and RMS cell lines did not alter the 3D collagen-induced apoptotic response, although verteporfin treatment caused a general reduction of proliferation (Supplementary Fig. 2C,D). Altogether, these data show that, unlike OS cells, collagen induces an apoptotic response in a group of RMS cell lines in a manner dependent on the 2-versus 3-dimensionality and microarchitecture of the matrix.

### DDR1 signalling mediates collagen-induced apoptosis

To understand the mechanisms behind the observed collagen-induced apoptotic responses, we compared the gene expression of the three collagen-apoptotic RMS cell lines (RH36, RMS, and RMSYM) and the three collagen-proliferative cell lines (RD, RH30, RUCH2). Pathway analysis of the differentially expressed genes revealed “muscle structure development”, “blood vessel development”, “VEGFA-VEGFR2 signalling pathway”, “cell morphogenesis involved in differentiation” and “cell-substrate adhesion” as the five most altered pathways between these two groups (Fig. 4A). Moreover, “positive regulation of cell death” and “apoptotic signalling pathway” also appeared significantly altered. Thus, we hypothesised that the expression of distinct collagen I receptors could result in the different response to collagen in distinct RMS cell lines as well as between RMS and OS cells. Consistently, the gene expression of collagen I receptors, linked to active tissue remodelling and cancer, including *DDR2, ITGA2, ITGA10, ITGA11*, and *ITGB1* was higher in OS compared to RMS, whilst *DDR1* expression was higher in RMS tumours (Fig. 4B). The expression of *DDR1* was also upregulated in RMS tumours compared to skeletal muscle and in more aggressive fusion+ compared to fusion-RMS tumours (Supplementary Fig. 3A and Fig. 4C). Moreover, *DDR1* was upregulated and *ITGB1* was downregulated in the collagen-apoptotic RH36, RMS, and RMSYM compared to the collagen-proliferative RD, RH30, and RUCH2 (Fig. 4D).

**Figure 4.**
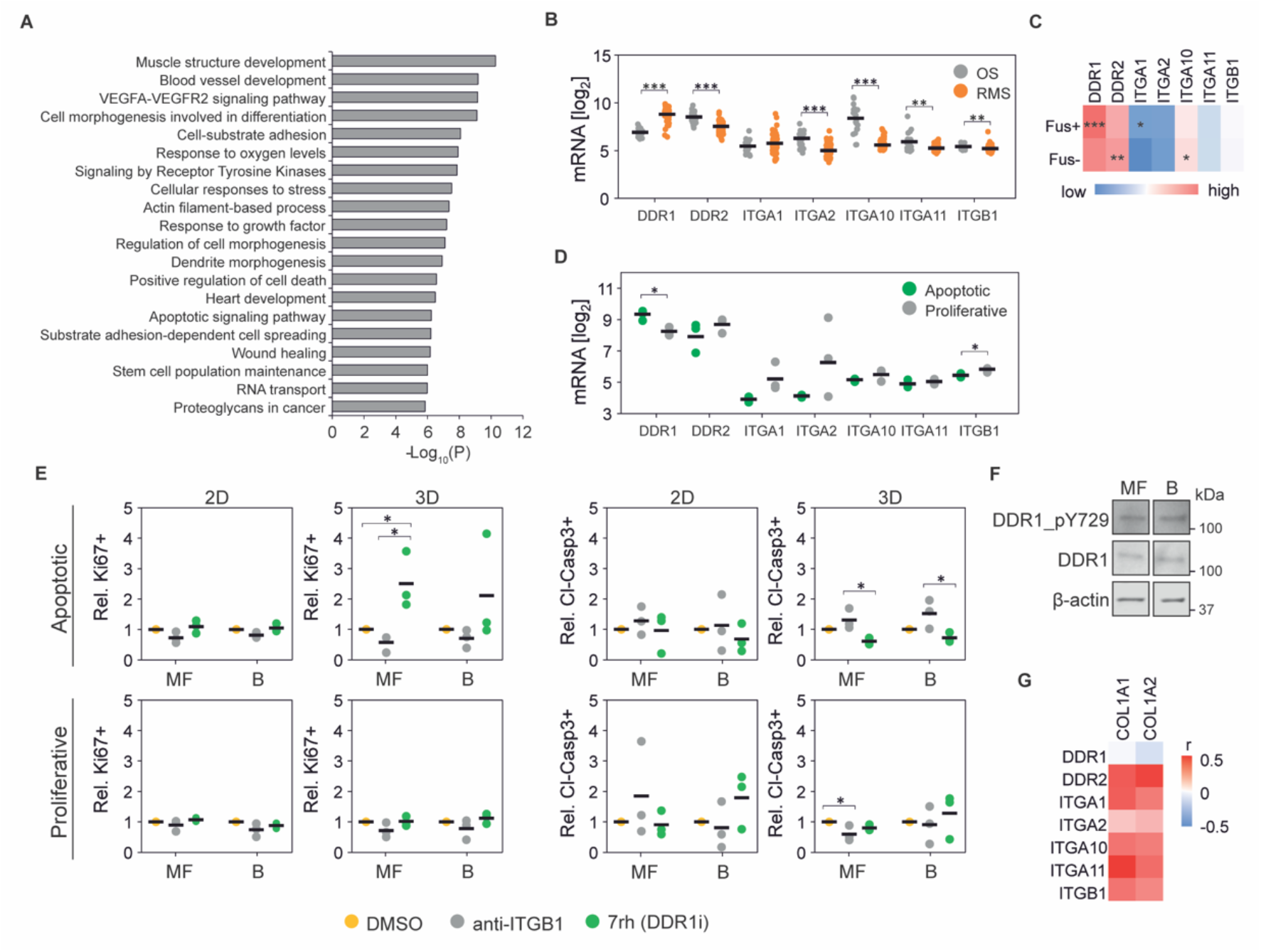
DDR1 signalling induces apoptosis in rhabdomyosarcoma cells. **A**, Most significantly altered pathways between apoptotic and proliferative cell lines at the gene level. **B**, Gene expression of collagen I receptors in osteosarcoma (OS; n=21; GSE87437) and rhabdomyosarcoma (RMS; n=58; GSE66533) tumours. **C**, Gene expression of collagen type I receptors in Fusion-positive (Fus+; n=45) and Fusion-negative (Fus-; n=56) rhabdomyosarcoma tumours. **D**, Gene expression of collagen I receptors in apoptotic and proliferative RMS cell lines. **E**, Proliferative and apoptotic response of cells upon treatment with integrin β1 blocking antibody (anti-ITGB1) or 7rh show increased percentage of Ki67+ and reduced cleaved-caspase 3+ cells in apoptotic cell lines treated with 7rh in 3D conditions (n= 3 per cell line). **F**, Protein expression of RMSYM cells seeded on MF- and B-collagen show no differences in DDR1 activation. **G**, Pearson ‘s correlation between collagen I and collagen I receptor gene expression in rhabdomyosarcoma tumours (n=101). Plots indicate individual data points and the average is shown as a bar. *** p < 0.001, ** p < 0.01, * p < 0.05.

To determine if DDR1 and integrin β1 functionally contribute to the collagen-dependent differences in RMS proliferation and apoptosis, we treated cells with an integrin β1 blocking antibody or the DDR1 inhibitor 7rh in 2D or 3D and in MF- or B-collagen. Notably, inhibition of DDR1 increased proliferation and reduced apoptosis in the apoptotic RH36, RMS, and RMSYM, in 3D especially MF-collagen (Fig. 4E). Overexpression of DDR1 in turn reduced proliferation and enhanced apoptosis of the RH30 cells, otherwise proliferative in 3D MF-collagen (Supplementary Fig 3B,C). In contrast, the integrin β1 inhibition tended to reduce proliferation and increase apoptosis (Fig. 4E). Subsequently, we sought to understand the reduced apoptotic response and enhanced proliferation observed in RMSYM cells on B-collagen (Fig 3B,F). In these cells, activation of DDR1, indicated by the levels of tyrosine 729 phosphorylation (pDDR1), was not affected by collagen microarchitecture (Fig. 4F). Moreover, RMSYM cell proliferation was strongly reduced and the apoptotic response enhanced by integrin β1 blockade in 3D B-collagen (Supplementary Fig. 3D,E). These results, together with the observed increase in integrin β1 activation in RMSYM cells on 2D B-collagen compared to MF-collagen (Fig. 2D,F), indicate that B-collagen-induced integrin β1 activation regulates cell survival and proliferation in these cells.

Finally, to understand the impact of DDR1 and collagen I on tumour development and aggressiveness in RMS patients, we compared the overall survival of patients with high and low *DDR1* gene-expressing tumours. Probability of survival analysis revealed that *DDR1* expression is not associated with patient prognosis (Supplementary Fig. 3F). However, gene expression correlation analysis indicated that the expression of all collagen I receptors except *DDR1* positively correlates with collagen I genes whilst *DDR1* expression correlates negatively, indicating that collagen-rich tumours generally express low *DDR1* (Fig. 4G). Altogether, these results show that DDR1 is a mediator of collagen-induced apoptosis of RMS cells and reveals reduction of DDR1 expression as a strategy to allow RMS cell growth in collagen-rich microenvironments.

### Matrix remodelling regulates collagen-induced apoptosis and DDR1 signalling

Single cells embedded within a 3D collagen matrix are exposed to a mechanically confining microenvironment. This confinement can be overcome by the cell-induced mechanical or proteolytic remodelling of the surrounding collagen. To investigate whether matrix remodelling is involved in collagen-induced RMS cell apoptosis, we first compared the mechanical remodelling of the apoptotic and proliferative RMS cell lines. Displacement of beads entrapped within MF-collagen matrices caused by collagen remodelling was similar in collagen-apoptotic and -proliferative cells, with the exception of RUCH2 presenting large displacements, indicating that the lack of mechanical remodelling alone does not necessarily cause apoptosis (Fig. 5A,B). Moreover, the expression of genes for matrix metalloproteinases (MMP) involved in collagen I proteolysis was, in general, lower in RMS than in OS tumours (Supplementary Fig. 4A). The expression of collagen I-degrading MMP genes was similar amongst the distinct RMS cell lines except RUCH2, which highly expressed *MMP1* (Supplementary Fig. 4B). To investigate the impact of collagen remodelling on proliferation and apoptosis, we inhibited MMPs with the broad-spectrum inhibitor GM6001, and mechanical remodelling with the myosin II inhibitor blebbistatin. As expected, the inhibition of myosin II or the MMPs, as well as their combined inhibition caused a significant reduction of Ki67 expression in cells embedded within 3D MF-collagen (Fig. 5C,D). However, the effect of these inhibitors on apoptosis was less pronounced, showing a significant increase only in blebbistatin-treated RMSYM (Fig. 5E,F).

**Figure 5.**
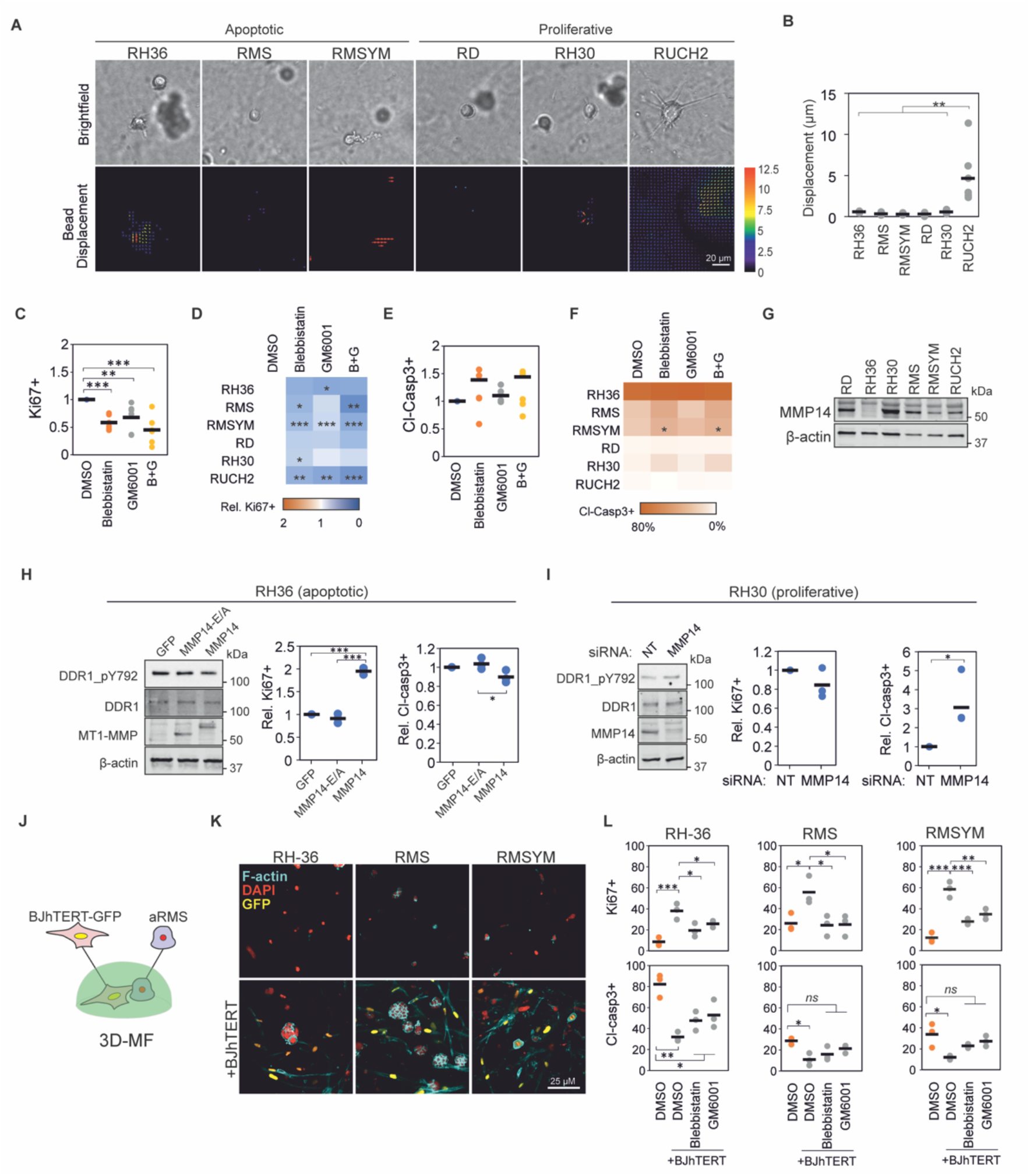
Matrix remodelling regulates DDR1 signalling and collagen-induced apoptosis. **A**, Representative examples of brightfield images (top) and corresponding bead displacement maps (bottom) through the first 4 h after embedding rhabdomyosarcoma cell lines in MF-collagen. Scale bar indicates 20 μm. **B**, Quantification of average bead displacement (n=6 cells). **C**, Quantification of relative Ki67+ cells of cells in 3D MF-collagen upon indicated treatments showing reduced Ki67+ cells compared to control. Each data point represents the average of a cell line (n=3 per cell line). **D**, Results from (**C**) represented per cell line. **E**, Quantification of relative cleaved caspase 3+ cells from (**C**). **F**, Results from (**D**) represented per cell line. **G**, Protein expression of MMP14 in each rhabdomyosarcoma cell line. **H**, Overexpression of wild-type MMP14 (MMP14) but not the catalytically inactive MMP14 mutant (MMP14-E/A) induces reduction of phospho-DDR1 without changes in total DDR1 in RH36 cells (left panel), increases the percentage of Ki67+ cells in 3D MF-collagen (central panel), and shows a non-significant tendency to reduction of percentage of cl-caspase 3+ (right panel) in cells embedded in 3D MF-collagen (n=3). **I**, Knockdown of MMP14 in RH30 cells increases phospho-DDR1 without changes in total DDR1 (left panel), reduces percentage of Ki67+ cells (central panel), and increases the percentage of cleaved-caspase 3+ cells (right panel) in cells embedded in 3D MF-collagen (n=3). **J**, Schematic representation of co-culture experiments of BJhTERT-GFP fibroblasts and apoptotic rhabdomyosarcoma cell lines (aRMS) embedded in 3D MF-collagen. **K**, Cells from (**J**) or in mono-culture after 4 days in culture. Scale bar indicates 25 μm. **L**, Percentage of Ki67+ (top) and cleaved-caspase 3+ cells (bottom) of cells from (**J**) after 24 h in culture with indicated treatments (n=3 per cell line). Plots indicate individual data points and the average is shown as a bar. *ns* (not significant) p > 0.05, *** p < 0.001, ** p < 0.01, * p < 0.05.

The activity of MMP14, central regulator of collagen remodelling and membrane receptor signalling, has been shown to regulate DDR1 signalling in breast cancer cells (*22*). Thus, we compared the MMP14 protein expression between the distinct RMS cell lines and investigated its function on regulating DDR1 signalling and apoptosis. Notably, the collagen-proliferative RD and RH30 express the highest MMP14 levels, whilst the apoptotic RH36 present the lowest MMP14 expression (Fig. 5G). In RH36 cells, overexpression of MMP14, but not a catalytically-inactive MMP14 mutant, increased proliferation and reduced apoptosis (Fig. 5H). Concurrently, knockdown of the high expression of MMP14 in RH30 cells enhanced apoptosis but had no significant effects on proliferation (Fig. 5I). Furthermore, overexpression of MMP14 caused a mild reduction and MMP14 knockdown an increase in pDDR1 without major changes in total DDR1 expression. These data indicate that MMP14 is a regulator to overcome collagen-induced apoptosis in RMS, which function can be partly attributed to changes in DDR1 signalling.

Finally, to investigate whether ECM remodelling induced by other cells of the TME is sufficient to promote cell survival on collagen-apoptotic RMS cell lines, we co-cultured the apoptotic RMS cell lines with actively collagen-remodelling fibroblasts (Fig. 5J). After 4 days in culture, RH36 and RMS cell lines formed spheroids, and RMSYM presented large cells similar to 2D conditions (Fig. 5K). Moreover, cell proliferation was enhanced and apoptosis reduced in the RMS cells in the co-cultures after 24 h incubation, an effect that was partially reduced upon myosin II and MMP inhibition (Fig. 5L). Altogether, these results indicate that matrix remodelling induces survival in collagen-apoptotic RMS cell lines, through mechanisms at least partially dependent on DDR1.

### 3D collagen-induced mechanical confinement synergises with DDR1 signalling to regulate apoptosis

Next, we sought to understand the effect of 3-dimensionality on collagen-induced apoptosis, which could indicate other mechanisms of matrix remodelling on RMS cell survival. First, to explore the mechanical stress experienced by cells, we compared the cellular and nuclear morphology of collagen-apoptotic RH36 cells on 2D and in 3D MF-collagen (*23*). After 3.5 hours in culture, prior to the initiation of the apoptotic response (Supplementary Fig. 5A-C), the cellular but not the nuclear volume was larger in 3D than in 2D conditions (Fig. 6A-C). Interestingly, nuclei in 3D conditions presented higher curvature (Fig. 6D), a morphology observed in MMP-inhibited cells migrating through the small pores of collagen matrices (*24*). This 3D collagen-induced nuclear deformation suggests that these cells are under anisotropic mechanical confinement. To investigate whether changes in mechanical confinement regulate 3D collagen-induced apoptosis, we decreased cell confinement by subjecting cells to hyposmotic conditions during and after collagen gel formation. Hyposmotic pressure reduced the nuclear curvature (Fig. 6E,F), enhanced proliferation (Fig. 6G,H), and reduced apoptosis in collagen-apoptotic cells (Fig. 6G,I and Supplementary Fig. 5D). To explore if the hyposmotic pressure-induced cell survival derived from reduced DDR1 activation, we compared the levels of pDDR1 in RH36 under isosmotic, hyperosmotic, and hyposmotic conditions. The levels of pDDR1 increased in hyposmotic pressure and decreased in hyperosmotic pressure, indicating that the effect of osmotic pressure on survival is not caused by the reduction of pDDR1 (Fig. 6J). Yet, the osmolarity-induced changes in pDDR1 suggest a mechanical regulation of DDR1 activity.

**Figure 6.**
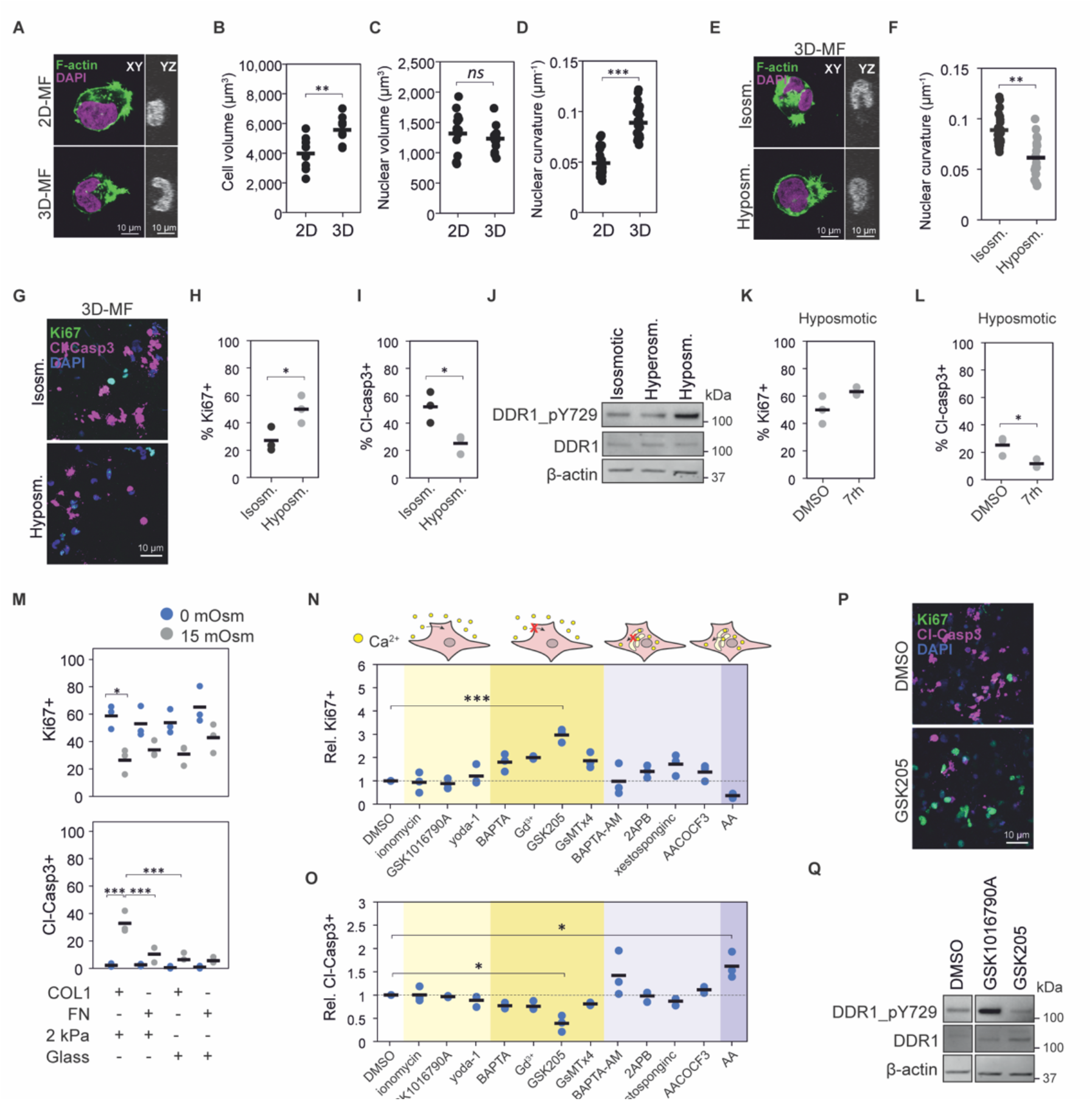
Mechanical confinement regulates apoptosis through DDR1-dependent and -independent mechanisms. **A**, Representative images of RH36 cells on 2D MF-collagen or embedded in 3D MF-collagen for 3.5 h. Scale bars indicate 10 μm. **B, C, D**, Quantification of cell volume (**B**; n=9 cells), nuclear volume (**C**; n≥13 cells), and nuclear curvature (**D**; n≥22 cells), based on F-actin and DAPI staining of cells from (**A**). **E**, Representative images of RH36 cells in 3D MF-collagen in isosmotic or hyposmotic conditions for 3.5 h. Scale bars indicate 10 μm. **F**, Nuclear curvature quantification of cells in (**E**). **G**, Representative images of Ki67+ and cleaved caspase 3+ RH36 cells embedded in 3D MF-collagen for 24 h in isosmotic or hyposmotic conditions. **H, I**, Quantification of Ki67+ (**H**) and cleaved caspase 3+ (**I**) cells from (**G**). **J**, Changes in osmotic pressure affect the levels of DDR1 phosphorylation in RH36 cells seeded on 2D MF-collagen for 6 h. **K, L**, Quantification of Ki67+ (**K**) and cleaved caspase 3+ (**L**) RH36 cells embedded in 3D MF-collagen in hyposmotic conditions for 24 h (n=3). **M**, Quantification of Ki67+ and cleaved caspase 3+ RH36 cells seeded on 2 kPa polyacrylamide gels or glass coverslips functionalised with collagen I or fibronectin in isosmotic or hyperosmotic conditions. **N, O**, Relative proportion of Ki67+ cells (**N**) and cleaved-caspase 3+ cells (**O**) of RH36 cells embedded in 3D MF-collagen for 24 h in the indicated treatments (treatments grouped by their effect on extracellular and intracellular calcium) showing significant increase in Ki67+ and decreased cleaved-caspase 3+ cells in cells treated with the TRPV4 inhibitor GSK205 (n=3). **P**, Representative images of RH36 cells in (**N**) and (**O**) showing decreased cleaved caspase 3+ and increased Ki67+ cells in GSK205-treated cells. **Q**, Effects of treatments with TRPV4 agonist GSK1016790A and antagonist GSK205 on DDR1 phosphorylation in RH36 cells on 2D MF-collagen. *** p < 0.001, ** p < 0.01, * p < 0.05.

To investigate whether DDR1 activity and mechanical confinement synergise to induce apoptosis in RMS cells, we inhibited DDR1 in cells in 3D MF-collagen under hyposmotic conditions. Inhibition of DDR1 further reduced 3D collagen-induced apoptosis without significant changes in proliferation (Supplementary Fig. 5D and Fig. 6K,L). Furthermore, on 2D MF-collagen, DDR1 inhibition attenuated the apoptosis response caused by hyperosmotic pressure, indicating that mechanical confinement and DDR1 signalling synergise to induce apoptosis (Supplementary Fig. 5E,F). Subsequently, we investigated whether other means of mechanical confinement can induce apoptosis. Embedding cells in high density 2% agarose gels, which induces isotropic mechanical confinement, or low density 0.2% agarose revealed that cell proliferation was generally lower in 2% agarose in all cell lines, but apoptosis was only induced in collagen-apoptotic cell lines (Supplementary Fig. 5G).

Finally, to investigate whether cell adhesion and substrate stiffness can regulate the apoptotic response to mechanical confinement, we seeded cells on top of collagen I or fibronectin-functionalised soft 2 kPa polyacrylamide gels, where the cells do not spread, or very stiff glass, where cells adhere and spread efficiently (Supplementary Fig. 6A). Hyperosmotic pressure reduced proliferation and induced apoptosis in collagen-apoptotic cell lines adhered to 2 kPa but not to glass substrates (Supplementary Fig. 6B and Fig. 6M). Furthermore, in RH36, hyperosmotic pressure induced apoptosis in collagen-functionalised but not in fibronectin-functionalised 2 kPa gels (Fig. 6M). Altogether, these results indicate that DDR1 signalling and mechanical confinement synergise to regulate apoptosis, and that adhesion to stiff substrates can overcome these apoptotic signals.

### TRPV4 regulates DDR1 activation and 3D collagen-induced apoptosis

Collagen-induced confinement alters calcium signalling by modulating the activity of stretch-activated mechanosensitive channels at the plasma membrane or the endoplasmic reticulum (ER) (*23, 25*). Moreover, changes in cell volume caused by osmotic pressure or cell-substrate adhesion control the activity of these channels (*26, 27*). Additionally, calcium is a mediator of apoptosis (*28*). Thus, to assess if mechanosensitive calcium channels regulate the effects of confinement and osmotic pressure on apoptosis and DDR1 activity, we compared the intracellular calcium levels of RMS cells in 2D and 3D during the first 3 h of culture. On average, the total intracellular calcium was lower in apoptotic cell lines than in proliferative cell lines, especially in 3D (Supplementary Fig. 7A,B). Furthermore, apoptotic cells showed higher calcium levels in 2D than in 3D, whilst proliferative cells presented higher calcium levels in 3D (Supplementary Fig. 7C). However, most cell lines in 3D showed increased calcium levels after 3 h compared to 1 h in culture (Supplementary Fig. 7D), which can be the result of mechanosensitive channel activation caused by cell volume expansion or membrane deformations (Fig. 6B).

**Figure 7.**
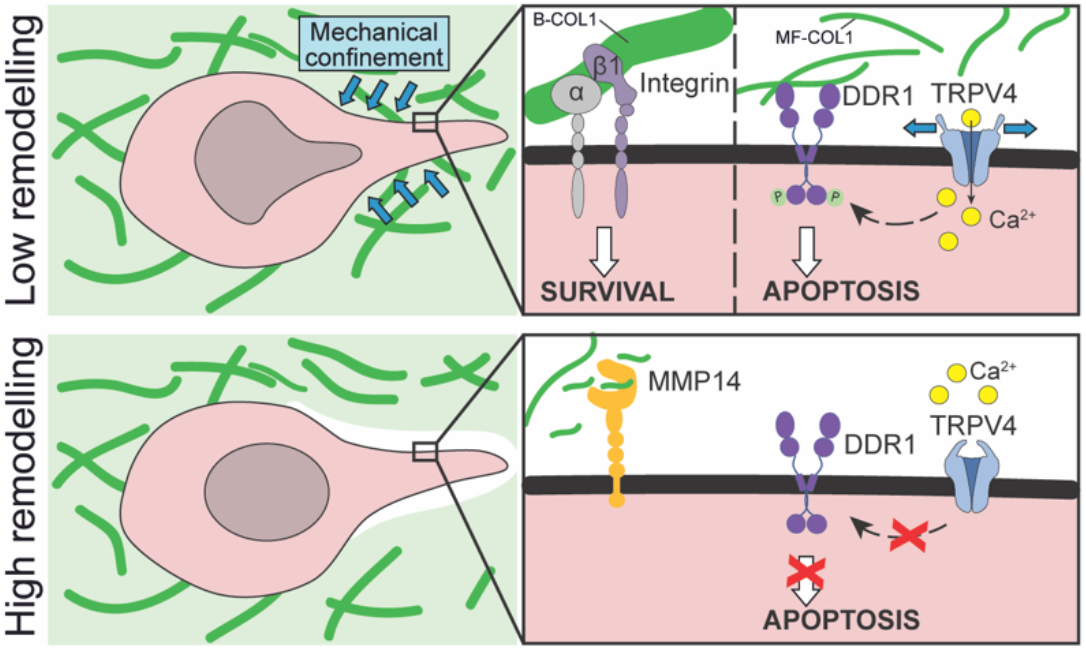
Proposed mechanism of collagen-induced apoptosis in rhabdomyosarcoma. In rhabdomyosarcoma cells with low collagen-remodelling capabilities, soft microfibrillar collagen causes apoptosis through biochemical and biomechanical signals. Biochemically, collagen induces an apoptotic response through the activation of DDR1. Biomechanically, the confinement imposed by the surrounding collagen induces a DDR1-independent apoptotic response but also activates the mechanosensitive ion channel TRPV4, which, in turn, enhances DDR1 activation leading to apoptosis. In contrast, stiff bundled collagen favours the activation of integrin β1 promoting cell survival. In cells with high remodelling capabilities, such as those highly expressing MMP14, there is reduced DDR1 signalling and confinement, favouring cell survival.

To investigate whether calcium signalling regulates the 3D collagen-induced apoptotic response, we activated or blocked extracellular calcium influx or calcium release from the ER with small molecule inhibitors, agonists, and calcium chelators. Extracellular calcium chelation with BAPTA and inhibition of mechanosensitive channels with gadolinium or the Piezo1, TRPC1, and TRPC6 inhibitor GsMTx4 lead to a non-significant trend for increased proliferation and decreased apoptosis (Fig. 6N,O). Instead, the selective antagonist of TRPV4 GSK205 significantly enhanced proliferation and reduced apoptosis (Fig. 6P). Neither the activation of membrane channels nor the blockage or stimulation of ER calcium release enhanced cell survival. Expression of TRPV4 was comparable across the cell lines except in RUCH2, which showed lower expression, indicating that TRPV4 is not the sole factor driving the apoptotic response to 3D collagen (Supplementary Fig. 7E). However, treatment with the TRPV4 agonist GSK1016790A increased and GSK205 decreased DDR1 phosphorylation (Fig. 6Q), consistent with the effects of osmotic pressure on DDR1 activity (Fig. 6J). Moreover, at the tumour level, *TRPV4* gene expression was lower in the more aggressive fusion+ than in fusion-tumours, opposite to the *DDR1* expression trend (Supplementary Fig. 7F). Altogether, these results show that, besides DDR1-independent effects of mechanical confinement on apoptosis, mechanical confinement also regulates DDR1 activity and apoptosis through TRPV4 in RMS cells.

## DISCUSSION

Expression of a dense fibrillar collagen-rich ECM in tumours is associated with aggressive cancer cell behaviour and unfavourable patient prognosis in various carcinoma types including breast, liver, pancreatic, ovarian, and lung cancer amongst others (*29*). Increased knowledge of the mechanisms leading to the ECM-promoted cancer aggressiveness has prompted the development of novel therapeutic strategies, currently under clinical trials (*6*). The impact of fibrillar collagens in sarcomas, which arise from cells of mesenchymal origin, is far less understood, likely due to their rareness. Thus, understanding the impact of fibrillar collagens in specific types of sarcoma is fundamental to evaluate their potential for therapeutic interventions and as diagnostic and prognostic biomarkers.

In this study, we investigated the impact of fibrillar collagens on RMS cell functions. To this end, we developed a method to modify collagen microarchitecture based on inducing high macromolecular crowding. This method, unlike other methods based on the control of temperature and pH (*17*), allowed us to recapitulate the vastly different collagen microarchitecture of distinct RMS-relevant tissues, whilst maintaining the temperature, pH, and osmotic pressure at physiological levels. For instance, using the high macromolecular crowding method, we obtained lung-like collagen structures, mimicking the microenvironment of the most common metastatic site in RMS (*18*). Normal lung is characterised by the presence of long collagen bundles and low collagen density. We show that both low collagen density and lung-like B-collagen favour RMS cell survival and proliferation, indicating that the lung provides an ideal milieu for the seeding of RMS metastasis.

Unlike in other major sarcoma and carcinoma types, using transcriptomic data of RMS tumours from the ITCC cohort, we demonstrate that the expression of fibrillar collagens and positive regulators of collagen biosynthesis and crosslinking was unexpectedly associated with better overall patient survival in RMS. Our results further indicate that RMS tumours express relatively high DDR1, which activity in corresponding cells was involved in collagen-induced apoptosis. In other types of normal and cancer cells, pro-apoptotic and anti-proliferative signals through DDR1 and integrin β1, respectively, have been reported (*22, 30*). By physically limiting cellular volume expansion, collagen is also able to inhibit cell growth, which effect is tightly linked to low collagen remodelling activity in cells (*31*). Since RMS cells are characterised by low collagen-remodelling matrix metalloproteinases, including the cell-surface collagenase MMP14 (*10*), the low collagen-remodelling activity and high DDR1 expression can render RMS cells especially susceptible to collagen-induced apoptosis. This suggests that the low collagen-remodelling activity and high DDR1 expression in RMS cells makes them susceptible to collagen-induced apoptosis. Such collagen-apoptotic response explains the unique association between high fibrillar collagen expression and favourable RMS patient outcome as well as the preference of RMS cells to metastasise to the lung and other low collagen content and confinement tissues.

Exploring the specific molecular mechanisms behind these observations, our results reveal that DDR1 activity is involved in an apoptotic response caused by 3D but not 2D collagen, especially in softer MF-collagen. However, the stiffer B-collagen favours RMS cell survival and proliferation through the activation of integrin β1, often associated with cancer cell survival (*32*), allowing cells to overcome the collagen-DDR1 pro-apoptotic signals. Thus, enhanced integrin β1 activation in lung-like B-collagen may promote RMS cell survival and proliferation in the lung.

Amongst the sarcoma cell lines used in this study, collagen caused apoptosis in the cell lines expressing high DDR1, low integrin β1, and relatively low MMP14, suggesting that matrix remodelling is also a central factor in the collagen-induced apoptotic response consistent with previous reports in breast cancer (*33*). Indeed, we found that overexpressing MMP14 or co-culturing apoptotic cell lines with highly remodelling fibroblasts, enhances the survival of these cells. These results suggest that non-malignant cells of the TME aid RMS cells to allow their proliferation through collagen remodelling, like previously reported in carcinomas (*4*). Expression analysis of these proteins in primary and distinct metastatic RMS tumours at various sites will be of interest to determine whether these factors (i.e. DDR1, integrin β1, and matrix remodelling through MMPs) are predictive of preferential metastatic sites.

Finally, we demonstrate that DDR1 activation and mechanical confinement synergise to induce apoptosis. The collagen-induced apoptosis is specifically induced in 3D but not in 2D cultures due to the mechanical confinement. The mechanical confinement of cells induces adhesion-independent apoptosis through YAP inhibition and by rupturing the nuclear envelope of migrating cells (*34, 35*). However, the collagen-induced apoptotic response in RMS cells is both 3D and DDR1-mediated adhesion-dependent. This synergy results from both DDR1-independent pro-apoptotic signals and the activation of DDR1 by the mechanosensitive ion channel TRPV4, which is activated by mechanical confinement as a result of cell membrane stretching (*31, 36*). Consequently, the mechanical confinement in 3D cultures results in enhanced pro-apoptotic signals leading to RMS cell apoptosis.

Collectively, our current findings reveal a mechanism of collagen-induced apoptosis caused by mechanical confinement and DDR1 signalling, which may explain the association between fibrillar collagen-related gene expression and favourable patient prognosis as well as the preference of RMS cells to metastasise in specific tissues (Fig. 7). Moreover, we describe molecular mechanisms that help explain the differential behaviour of cells to distinct culture dimensionality and microenvironmental collagen architecture, highlighting the importance of choosing cancer-type specific in vitro models.

## MATERIALS AND METHODS

### Bioinformatic analysis

Gene expression data of human tumour and normal tissues were obtained from publicly available datasets including, the Gene Expression Omnibus (GEO) GSE66533, GSE87437, GSE141690. Gene expression and sarcoma patient clinical data was obtained from the Innovative Therapies for Children with Cancer consortium cohort (rhabdomyosarcoma), the GEO dataset GSE21257 (osteosarcoma), and the adult sarcoma cohort from The Cancer Genome Atlas program TCGA-SARC. The RNA raw data were normalized and compared using the Affymetrix Expression Console Software (Thermofisher). Pathway analysis of differentially expressed genes was conducted by Metascape (*37*).

### Picrosirius Red staining

Tissues were deparaffinized and rehydrated through graded ethanol series. Slides were incubated for 1 h at RT in picrosirius red (PSR) staining solution (1% (w/v) Direct Red 80 (2610-10-8; Sigma-Aldrich) in saturated picric acid (P6744-1GA; Sigma-Aldrich). Slides were washed twice with 0.5% acetic acid, dehydrated in graded ethanol series, cleared in Tissue Clear, and mounted with Pertex (Histolab, Cat # 00811). Slides were imaged with the Zeiss LSM800-Airy confocal microscope using the 561 nm laser. Image quantification was performed with the macro TWOMBLI of Fiji-ImageJ (*38*) and the ridge detection plugin.

### Cell culture

RH30 and RMSYM cells were maintained in RPMI and RD, RH36, RMS, RUCH2, and BJ-hTERT human fibroblasts in DMEM. All culture media were supplemented with 10% FBS and 100 mg/ml penicillin/streptomycin. The cells were checked for mycoplasma regularly.

### Collagen-based *in vitro* models

Rat tail collagen type I (Sigma) was dissolved in 0.4% acetic acid to 4.5 mg/ml stock and diluted to 2.25 mg/ml in 2x MEM. The pH was then adjusted to 7.5 with sodium hydroxide. Subsequently, solutions containing 200% fraction volume of occupancy (calculated based on PEG hydrodynamic radii (*39*)) of PEG at molecular weights of 8, 35, 300, or 1000 kDa (Sigma) in PBS were mixed 1:1 with the collagen solution. To control for effects of osmotic pressure, controls with equivalent osmotic pressure but low macromolecular crowding were prepared with PEG 400 Da. Collagen-PEG mixtures were then transferred onto cell culture plates with or without cells for 2D cultures or with 150 cells/μl for 3D cultures and incubated for 1 h at 37ºC. Prior to addition of cell culture medium, collagen gels were washed with 2X with PBS to remove soluble PEG. Imaging of collagen was performed by reflectance confocal imaging with a Zeiss LSM800 confocal microscope. Image quantification was performed with the macro TWOMBLI of Fiji-ImageJ and the ridge detection plugin.

### Mechanical characterization of collagen hydrogels

The Young ‘s modulus of collagen matrices was determined by AFM using a Bioscope catalyst AFM (Bruker) integrated with an inverted Axio Observer (Zeiss). Measurements were performed in contact mode while matrices were submerged in PBS. Samples were probed with a 4.5 μm diameter borosilicate glass sphere attached to a silicon nitride cantilever (Novascan Tech). Prior to each set of measurements, thermal calibration was performed to determine the spring constant of the cantilever (∼0.07 N/m). For analysis, the curves were fitted to the Hertz spherical indentation model, and Poisson ‘s ratio was set to 0.5. Young ‘s moduli were calculated with the NanoScope 8.15 software.

### Live cell imaging

For apoptosis dynamics quantification, the medium of collagen-embedded cells was exchanged by phenol red-free medium containing 15 mM HEPES (Sigma) and 2 μM CellEvent™ Caspase-3/7 Green (Invitrogen). Cells were imaged (brightfield and green channels) every 4 h for a total of 72 h. For bead displacement analysis, FluoSpheres™ Carboxylate-Modified Microspheres, 0.2 μm, red fluorescent (580/605), 2% solids (Thermo Fisher) were pre-diluted 1:10 in 1% FBS and vortexed. Of this, 4 μl were added per 100 μl of collagen mixture before 1 h incubation at 37 ºC for the collagen solution to form a gel. The final collagen matrices were imaged at 3 and 4 h in culture. Images were analysed using ImageJ with Particle Image Velocimetry plugins. For mathematical analysis, the resulting vector sizes of bead movement within a 48 × 48 μm square around the cell were averaged. For calcium measurements, the Fluo-4 Calcium Imaging Kit (ThermoFisher) was used. The kit Component A was used in 50:50 proportion with phenol-red free medium, and cells were preincubated for 1h before trypsinisation and transfer into or onto collagen matrices. Fresh medium/calcium detector mix was added and the cells were cultured for up to 5 h for imaging. All live cell imaging was conducted in a Cytation 5 imaging reader at +37 °C and 5% CO2 (BioTekTM CYT5MPV).

### Osmotic pressure experiments

Hyposmotic treatment of 3D cell cultures, 2.25 mg/ml collagen solution adjusted at pH 7.5 containing the desired cell number were dissolved with the same volume of a 60% deionised water 40% PBS solution. After 1 h incubation at 37ºC, media were prepared with either 30% deionised water or 30% PBS (control). Hyperosmotic treatment of cells was performed by adding PEG 400 Da to culture media after cell embedding or seeding.

### Antibodies

The antibodies used were the following: primary antibodies for immunofluorescence were Ki67 (8D5; #9449; Cell Signaling Technology; 1:800), active integrin β1 (12G10; ab30394; Abcam; 1:400), YAP/TAZ (63.7; sc-101199; Santa Cruz Biotechnology; 1:200), cleaved-caspase 3 (Asp175; #9664 Cell Signaling Technology; 1:800). Secondary andibodies for immunofluorescence were goat anti-rabbit Alexa Fluor Plus 555 (#A32732, ThermoFisher Scientific; 1:1000) and goat anti-mouse Alexa Fluor Plus 499 (#A32731, ThermoFisher Scientific; 1:1000). Primary antibodies used for western blot were pDDR1 (Tyr792; #11994; Cell Signaling Technolgy; 1:500), DDR1 (D1G6; #5583; Cell Signaling Technolgy; 1:1000), b-actin (C-4; sc-47778; Santa Cruz Biotechnology; 1:2000), MMP14 (clone LEM-2/15.8; MAB3328; Merk; 1:2000), TRPV4 (ab39260; Abcam; 1:750). Secondary antibodies for western blot used were goat anti-mouse Immunoglobulins/HRP (P044701-2; Dako), goat anti-rabbit Immunoglobulins/HRP (P044801-2; Dako), IRDye® 800CW goat anti-rabbit IgG Secondary Antibody (926-32211; Li-COR), and IRDye® 680RD goat anti-Mouse IgG Secondary Antibody (926-68070; Li-COR).

### Immunofluorescence

For 2D immunofluorescence, cells were fixed with 4% PFA for 20 min at RT, blocked with 5% BSA (Biowest) 0.3% Triton-X (Sigma) in PBS for 30 min at RT and incubated with primary antibody in blocking buffer for 1 h at RT. Cells were incubated with Alexa Fluor secondary antibodies (Thermo Scientific) in blocking buffer for 30 min at RT and mounted with VECTASHIELD Antifade Mounting Medium with DAPI (Vector Laboratories). For 3D matrices, cells were fixed with 4% PFA for 1 h at RT followed by a 45 s post-fixation step with ice-cold 1:1 mixture of acetone-methanol. Matrices were then blocked with blocking buffer (15% FBS – 0.3% Triton-X in PBS) for 2 h at RT and incubated with primary antibody in blocking buffer for 4 h at RT. Multiple washing steps with 0.45% Triton-X in PBS were performed and matrices were incubated with Alexa Fluor secondary antibodies and phalloidin in blocking buffer for 2 h at RT. For actin staining, phalloidin Alexa Fluor 488 or 647 (ThermoFisher) was used together with the secondary antibodies. After several washes with 0.45% Triton-X in PBS and 1X with PBS, matrices were mounted on an object slide with VECTASHIELD Antifade Mounting Medium with DAPI. Confocal images of immunofluorescence stainings were obtained using a Zeiss LSM800 confocal microscope. Imagen analysis was performed on Fiji-ImageJ.

### Immunoblot

For cells cultured in 3D, collagen droplets were first digested with 1X collagenase/hyaluronidase (StemCell Technologies) to harvest cells. Then the cell pellets or 2D cultured cells were lysed with RIPA lysis buffer (50 mM Tris–HCl pH 7.4, 150 mM NaCl, 1% Igepal CA-630, 0.5% sodium deoxycholate) supplemented with 5 mM EDTA, protease inhibitor (cOmplete ULTRA tablet, Sigma), and phosphatase inhibitor (PhosSTOP tablet, Sigma) on ice. Debris was removed from the supernatants by centrifugation at 21,130 g for 15 min at 4°C. We then measured protein concentrations with the Pierce BCA Assay (Thermo Scientific). Lysates were mixed with 5X sample buffer (0.3 M Tris–HCl pH 6.8, 50% glycerol, 10% sodium dodecyl sulfate, 0.05% bromophenol blue) containing 0.5 M dithiothreitol and heat-denatured at 95°C for 10 min before separation in 4–20% Mini-PROTEAN TGX Precast Protein Gels (BioRad) and transferred to Trans-Blot Turbo Mini Nitrocellulose Transfer Packs (Bio-Rad). Membranes were blocked for 45 min with 5% milk (Cell Signaling Technologies) or 3% fish gelatin (Sigma) in Tris-buffered saline (TBS; 10 mM Tris, pH 7.6, 150 mM NaCl) and probed with primary antibody in TBS 0.1% Tween-20 (TBS-T) with 5% milk or 3% fish gelatin at the recommended dilutions at 4°C overnight.

Membranes were incubated with horseradish peroxidase (HRP)-conjugated secondary antibodies (Dako) or with IRDye Subclass-Specific Antibodies (LI-COR Biosciences) diluted in TBS-T for 1 h at RT, and the signal was detected using ECL chemiluminescent detection reagent (GE Healthcare) and visualized using ChemiDoc Imaging System (Bio-Rad) or using Odyssey Imaging System (LI-COR Biosciences).

### cDNA overexpression and siRNA knockdown

For protein overexpression, the expression constructs encoding wild-type MMP14 catalytically inactive MMP14 bearing an Ala substitution at Glu240 (referred to as MT1-E/A) or pCMV-DDR1-t2-HA (HG10730-CY; Sino Biological) were used (*40*). Briefly, 1 μg DNA was incubated with 5 μl FUGENE (Promega) and 100 μl optiMEM for 15 min. Then, 900 μl medium was added and the final mixture incubated with the cells for 48 h.

For knockdown, siRNA against human MMP14 (L-004145-00-0005; Dharmacon), DDR1 (L-003111-00-0005; Dharmacon), or non-targeting control siRNA (QIAGEN: SI03650318) were used. Briefly, 150 μl optiMEM and 4.5 μl lipofectamine were mixed and incubated for 5 min and a mixture of 150 μl optiMEM and siRNA were added and incubated for 20 min. Then, they mixtures were added to 1.2 mL serum-free medium. After 24 h in culture, the medium was exchanged to complete medium and incubated for another 24 h.

### ECM-coated substrates

To prepare polyacrylamide gels, glass coverslips were washed twice with 70% ethanol, activated with 0.1 M NaOH, and silanized with (3-aminopropyl)trimethoxysilane (Sigma). The glass coverslips were then treated with 0.5% glutaraldehyde (Sigma) for 30 min and washed extensively with sterile MilliQ water. Mixtures of MilliQ water, acrylamide monomers (Sigma), and crosslinker N,N-methylene-bis-acrylamide (Sigma) were prepared according to previously determined formulations to obtain 2 kPa gels (*41*). For the polymerization reaction, 5 μl of 10% ammonium persulfate (Sigma) and 0.75 μl N,N,N′,N′-tetramethylethylenediamine (Sigma) were added into 0.5 ml mixtures. To functionalize gels with ECM proteins, gels were first treated with 1 mg/ml N-sulfosuccinimidyl-6-(4′-azido-2′-nitrophenylamino) hexanoate (Sigma), which was activated by ultraviolet (UV) light. Finally, gels or silanized glass coverslips were incubated with 10 μg/ml collagen I from rat tail (Sigma) or human fibronectin (Sigma) for 3 h at room temperature (RT) and UV-sterilized prior to cell seeding.

### Pharmacological and small-molecule inhibitors and agonists

The pharmacological and small-molecule inhibitors and agonists used were the following: 7rh (2.5μM; Sigma) for DDR1 inhibition, rat anti-human CD29 (1:100; BD Bioscience) to block integrin β1, blebbistatin (1 μM; Enzo Life Sciences) for myosin II inhibition, GM6001 (10 μM; Calbiochem) as broad-spectrum MMP inhibitor, L-mimosine (400 μM; Santa Cruz Biotechnology) to arrest the cell cycle at G1, S-Trityl-L-Cysteine (STLC; 20 μM; Santa Cruz Biotechnology) to arrest at mitosis, ionomycin (10 μM; MedChemExpress) to increase cell membrane permeability to calcium, gadolinium chloride (10 μM; MedChemExpress) to inhibit mechanosensitive calcium channels, BAPTA (10μM; MedChemExpress) to chelate extracellular calcium, BAPTA-AM (10 μM; MedChemExpress) to chelate intracellular calcium, arachidonic acid (70 μM; MedChemExpress) to mimic signalling downstream of nuclear mechanosensing, AACOCF3 (28 μM; Tocris) to inhibit signalling downstream of nuclear mechanosensing, 2-Aminoethoxydiphenylborane (20 μM; MedChemExpress) as IP3 receptor agonist, Xestospongin c (10 μM; Tocris) as IP3 receptor agonist, GSMTx4 (10 μM; MedChemExpress) as Piezo 1 inhibitor, Yoda1 (6 μM; MedChemExpress) as Piezo 1 agonist, GSK205 (10 μM; MedChemExpress) as TRPV4 inhibitor, GSK1016790A (50 μM; MedChemExpress) as TRPV4 agonist.

### Statistical analysis

The overall survival (OS) probabilities were estimated and presented by Kaplan–Meier survival curves. The number of technical and biological repeats is reported in each figure caption. When possible, single biological repeat data points are provided in the graphs. Statistical significances were determined using two-tailed Student ‘s t tests (to compare two samples), one-way analysis of variance with Tukey ‘s multiple comparison test (to compare multiple samples) or log-rank test (to compare patient survival). For correlation analyses, Pearson correlation coefficient was used. P values less than 0.05 were considered statistically significant. P-values are depicted as *p < 0.05, **p < 0.01, ***p < 0.001. Microsoft Excel and Prism v8.0 software (GraphPad) were used for statistical analyses.

## ACKNOWLEDGEMENTS

We thank the Biomedicum Imaging Core at Karolinska Institutet for imaging facilities. We also thank David Lane and Marie Arsenian-Henriksson for the Cytation 5 microscope. We thank Raushan T. Kurmasheva for facilitating the rhabdomyosarcoma cell line RH36 gene expression data.

## Author contributions

J.G.-M. conceived the project. J.G.-M., K.M.K., B.R., L.M.-G., M.C.-N., conducted the *in vitro* experiments and analysed the data. G.K., J.W.C., and M.E. provided human tissue samples and gene expression data. J.G.-M., A.B., and C.dL. assisted and performed AFM measurements. J.G.-M., K.M.K., B.R., L.M.-G., M.C.-N., M.E., J.W.C., and K.L. interpreted the data. J.G.-M. wrote the manuscript with contributions from all authors.

## Data and materials availability

All data needed to evaluate the conclusions in the paper are present in the paper and/or the Supplementary Materials. Additional data related to this paper may be requested from the authors.

