## Supplementary Fig. for "Mechanical confinement and DDR1 signalling synergise to regulate collagen-induced apoptosis in rhabdomyosarcoma cells"

### Supplementary Fig. 1

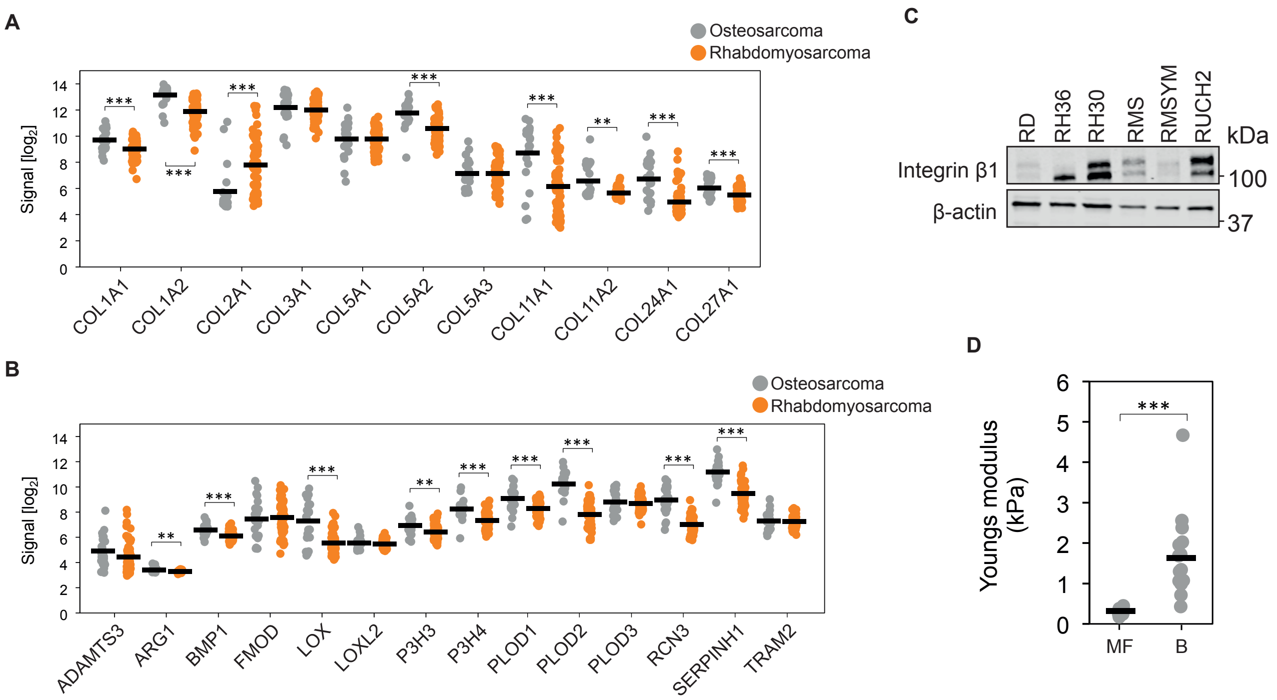

### Supplementary Fig. 2

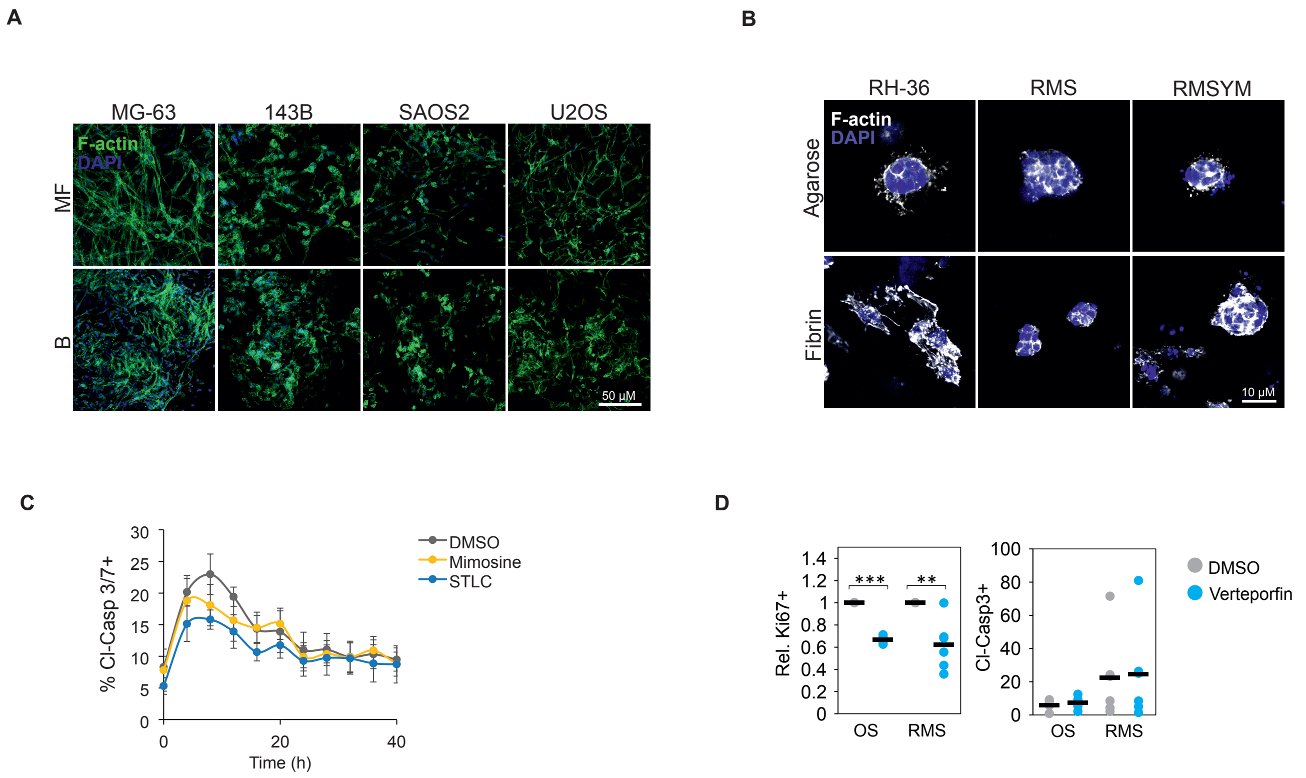

### Supplementary Fig. 3

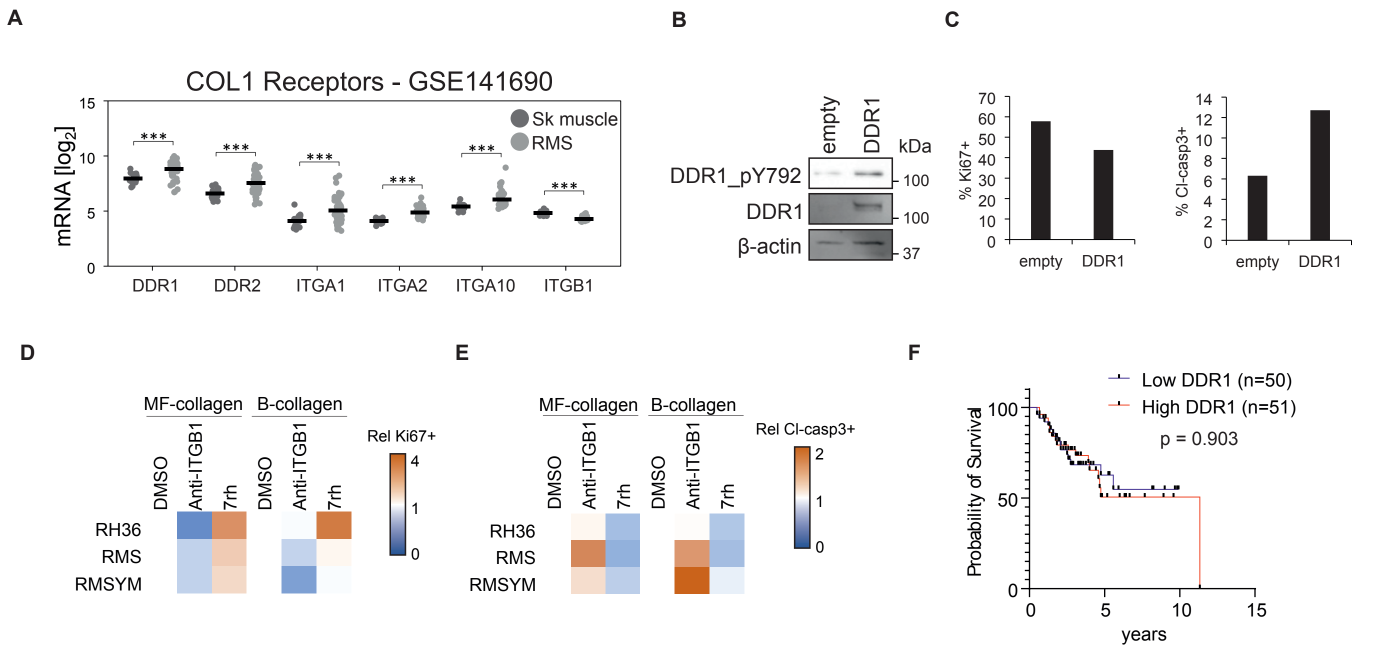

### Supplementary Fig. 4

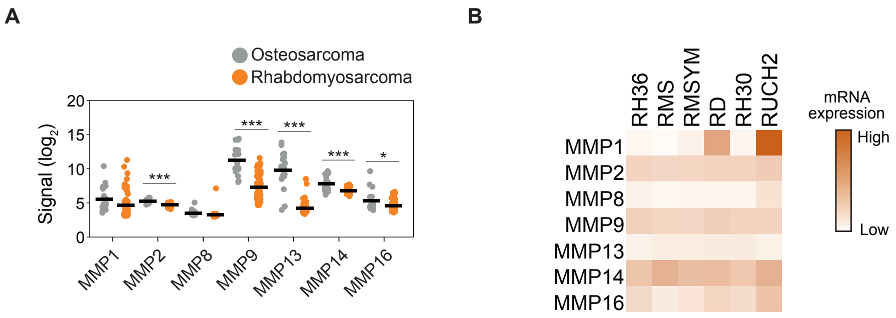

### Supplementary Fig. 5

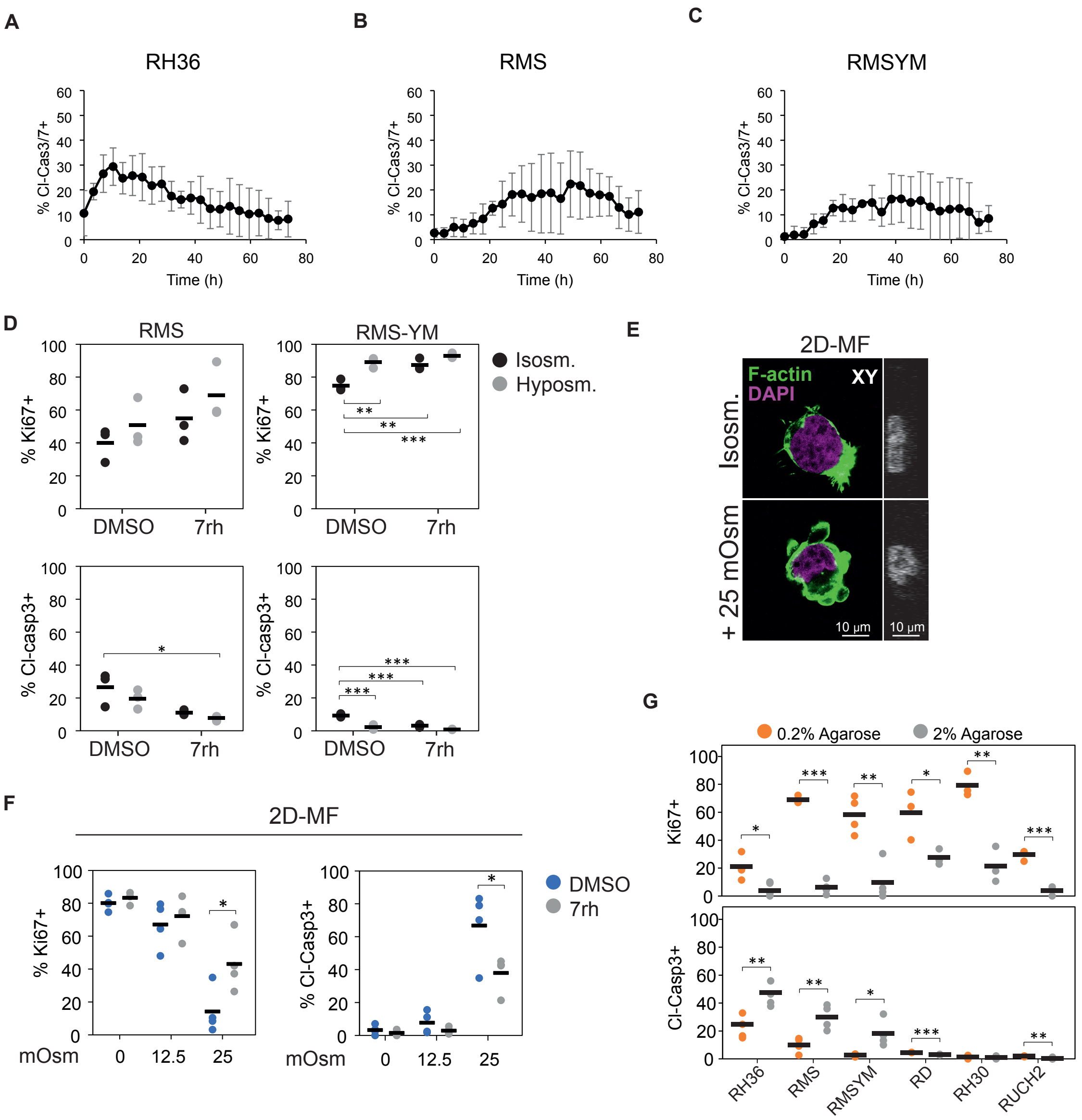

### Supplementary Fig. 6

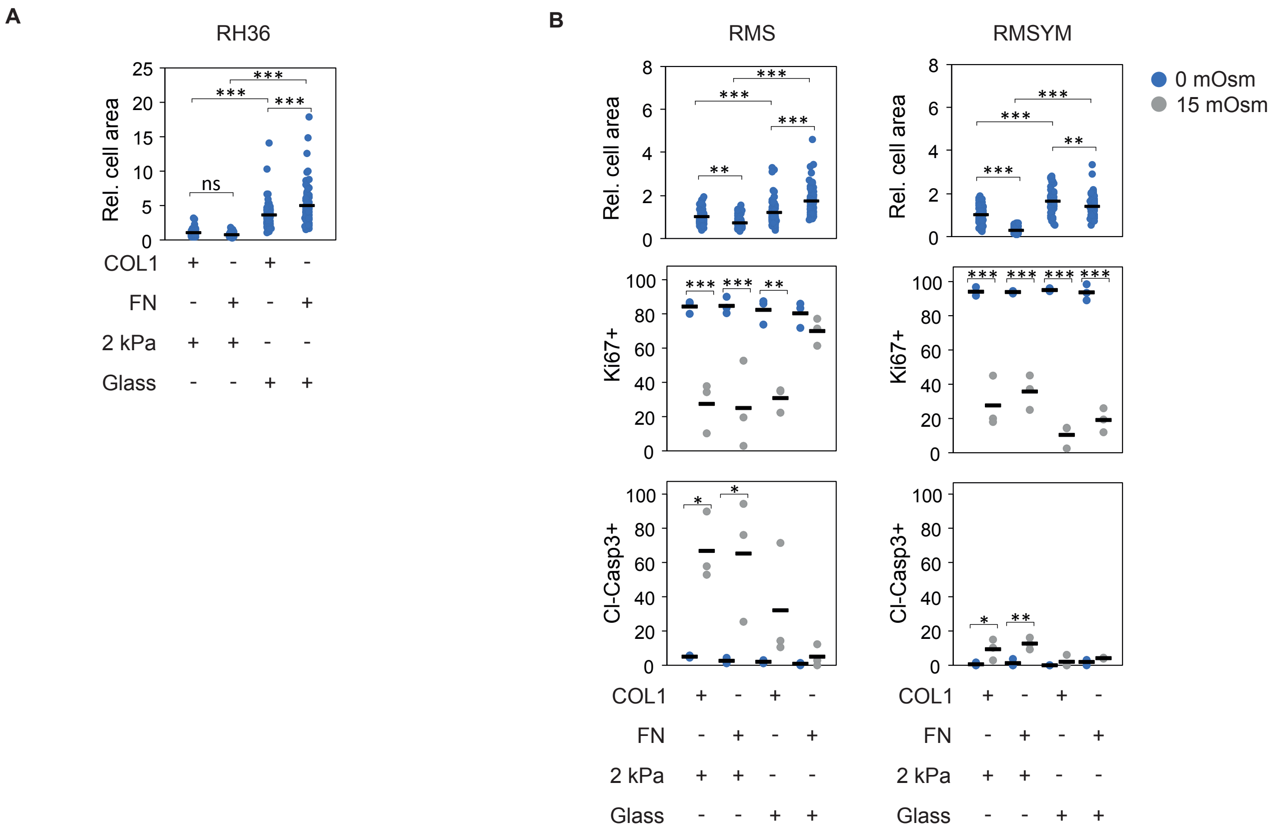

### Supplementary Fig. 7

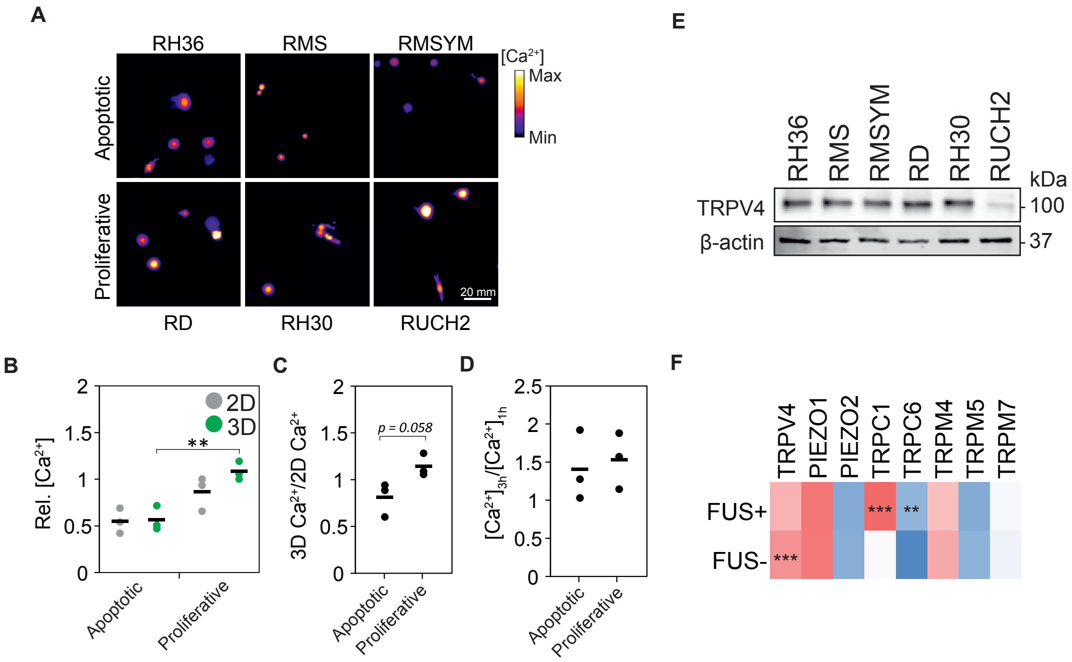

#### SUPPLEMENTARY FIGURE LEGENDS

**Supplementary Figure 1. Expression of collagen-related gene and integrin β1 protein in RMS and stiffness of collagen matrices with distinct microarchitecture. A**, **B**, Gene expression of fibrillar collagens (**A**) and collagen biosynthesis-related genes (**B**) in osteosarcoma (OS; n=21; GSE87437) and rhabdomyosarcoma (RMS; n=58; GSE66533) tumours. **C**, Integrin β1 protein expression in RMS cell lines. **D**, Young's moduli of microfibrillar-collagen (MF; n=17 spots) and bundle-collagen (B; n=16 spots). Each data point is shown, bars indicate average Young's moduli. *** $p < 0.001$, ** $p < 0.01$.

**Supplementary Figure 2. 3D collagen matrices induce apoptosis in RMS cell lines independently of cell cycle and YAP activation. A**, Representative images of OS cell lines embedded in microfibrillar (MF)- or bundled (B)-collagen for 4 days showing similar cell density in both matrices. Scale bar indicates 50 μm. **B**, Collagen-apoptotic RMS cell lines form spheroids in soft agarose and fibrin matrices after 4 days. Scale bar indicates 10 μm. **C**, Blocking cell cycle progression does not alter apoptosis dynamics induced by 3D MF-collagen on RH36 cells (n=3). **D**, Treatment with the YAP-TEAD inhibitor verteporfin reduces proliferation but does not alter apoptosis in cells embedded in 3D MF-collagen for 24 h. Each data point indicates average (n=3) of a cell line (OS n=4 cell lines, RMS n=6 cell lines). *** $p < 0.001$, ** $p < 0.01$.

**Supplementary Figure 3. Inhibiting DDR1 reduces and blocking integrin β1 enhances the apoptotic response to 3D collagen in RMS cells. A**, Comparison of collagen I receptor gene expression between skeletal muscle (Sk muscle; n=9) and rhabdomyosarcoma (RMS) tumours (n=66) showing a general increase except for ITGB1 in tumour tissues. **B**, Western blot analysis showing the efficiency of DDR1 overexpression in RH30 cells. **C**, Effect of DDR1 overexpression on RH30 cell proliferation (Ki67+) and apoptosis (Cleaved-caspase 3+) after 24 h embedding in 3D microfibrillar (MF)-collagen (n=3 technical repeats). **D**, **E**, Variation in cell proliferation (**D**) and apoptosis (**E**) in collagen-apoptotic RMS cell lines embedded in 3D MF-collagen for 24 h upon the indicated treatment showing opposite effects of DDR1 and integrin β1 inhibition (n=3 biological repeats). **F**, Kaplan-Meier survival curves of RMS tumours expressing low (n=50) or high (n=51) DDR1 (from the ITCC cohort) showing no difference between these groups. *** $p < 0.001$.

**Supplementary Figure 4. Matrix metalloproteinase gene expression is relatively low in RMS cells. A**, Comparison of collagen-proteolytic matrix metalloproteinase (MMP) gene expression between OS (n=21; GSE87437) and RMS (n=58; GSE66533) tumours. **B**, Gene expression of collagen-proteolytic MMPs in RMS cell lines.

**Supplementary Figure 5. Mechanical confinement synergises with DDR1 signalling to induce apoptosis in RMS cells. A-C**, Dynamics of apoptosis induced by MF-collagen in RH36 (**A**), RMS (**B**), and RMSYM (**C**) cells (n=5 technical repeats). **D**, Hyposmotic pressure and DDR1 inhibition synergise to enhance proliferation and reduce apoptosis in RMS and RMSYM cells embedded in 3D microfibrillar (MF)-collagen for 24 h (n=3). **E**, Representative images of RH36 cells on 2D MF-collagen gels subjected to distinct osmotic pressures showing membrane blebbing and changes in nuclear morphology in hyperosmotic conditions. Scale bars indicates 10 μm. **F**, Inhibition of DDR1 reduces apoptosis in RMS cells seeded on 2D MF-collagen subjected to hyperosmotic conditions (n=3). **G**, Apoptosis is enhanced in collagen-apoptotic RMS cells lines embedded in high density agarose compared to low density agarose hydrogels after 24 h (n=3). *** $p < 0.001$, ** $p < 0.01$, * $p < 0.05$.

**Supplementary Figure 6. Enhanced cell adhesion reduces hyperosmotic pressure-induced apoptosis. A**, Effect of adhesion to distinct substrates on RH36 cell spreading (n≥74 cells). **B**, Effect of cell adhesion to distinct substrates on RMS and RMSYM cell spreading (top panels; n≥75 cells), proliferation (middle panels; n=3 biological repeats), and apoptosis (bottom panels; n=3 biological repeats). *** $p < 0.001$, ** $p < 0.01$, * $p < 0.05$.

**Supplementary Figure 7. Intracellular calcium content and TRPV4 expression comparison in RMS cell lines. A**, Representative images of relative calcium content in RMS cell lines embedded in 3D microfibrillar (MF)-collagen for 3 h. Scale bar indicates 20 μm. **B-D**, Comparison of the relative calcium content (**B**), or the ratio of calcium content in cells on 2D and in 3D MF-collagen (**C**) after 3 h or the difference in calcium content between 1 h and 3h after embedding (**D**) in collagen-apoptotic and collagen-proliferative cell lines. Each data point indicates the average of one cell line and bars show the average of each group. **E**, Protein expression of TRPV4 in the distinct RMS cell lines. ** $p < 0.01$.
